# Evolutionary consequences of domestication on the selective effects of new amino acid changing mutations in canids

**DOI:** 10.1101/2024.11.13.623529

**Authors:** Carlos Eduardo G. Amorim, Chenlu Di, Meixi Lin, Clare Marsden, Christina A. Del Carpio, Jonathan C. Mah, Jacqueline Robinson, Bernard Y. Kim, Jazlyn A. Mooney, Omar E. Cornejo, Kirk E. Lohmueller

## Abstract

The domestication of wild canids led to dogs no longer living in the wild but instead residing alongside humans. Extreme changes in behavior and diet associated with domestication may have led to the relaxation of the selective pressure on traits that may be less important in the domesticated context. Thus, here we hypothesize that strongly deleterious mutations may have become less deleterious in domesticated populations. We test this hypothesis by estimating the distribution of fitness effects (DFE) for new amino acid changing mutations using whole-genome sequence data from 24 gray wolves and 61 breed dogs. We find that the DFE is strikingly similar across canids, with 26-28% of new amino acid changing mutations being neutral/nearly neutral (|*s|* < 1e-5), and 41-48% under strong purifying selection (|*s|* > 1e-2). Our results are robust to different model assumptions suggesting that the DFE is stable across short evolutionary timescales, even in the face of putative drastic changes in the selective pressure caused by artificial selection during domestication and breed formation. On par with previous works describing DFE evolution, our data indicate that the DFE of amino acid changing mutations depends more strongly on genome structure and organismal characteristics, and less so on shifting selective pressures or environmental factors. Given the constant DFE and previous data showing that genetic variants that differentiate wolf and dog populations are enriched in regulatory elements, we speculate that domestication may have had a larger impact on regulatory variation than on amino acid changing mutations.

**Significance Statement:** Domestication of dogs to live alongside humans resulted in a dramatic shift in the pressures of natural selection. Thus, comparing dogs and wolves offers a unique opportunity to assess how these shifts in selective pressures have impacted the fitness effects of individual mutations. In this project, we use patterns of genetic variation in dogs and wolves to estimate the distribution of fitness effects (DFE), or the proportions of amino acid changing mutations with varying fitness effects throughout the genome. Overall, we find that the DFE for amino acid changing mutations is similar between dogs and wolves. Even genes thought to be most affected by domestication show a similar DFE, suggesting that the DFE has remained stable over evolutionary time.

## Introduction

Domestication created radical phenotypic changes in many species and understanding the genetic and evolutionary basis of these changes is a major research objective (1–4). Studies on animal and plant domesticates have shown that these changes are accompanied by an increase in the number and frequency of deleterious genetic variants (5–7), an enrichment of identity-by-descent (IBD) segments, coupled with an excess of runs of homozygosity (ROH) (8, 9), the simplification of the genetic architecture of polygenic traits (7, 9), and an increase in the overall recombination rates (10, 11). What remains unclear is whether domestication has also altered the strength of natural selection on amino acid changing (or nonsynonymous) variants due to shifted selective pressures.

Artificial selection for desired traits during the process of domestication is thought to have led to the rapid increase in the frequency of alleles with large effects on the trait, triggering selective sweeps (7, 9, 12, 13), but may also have led to the relaxation of the selective pressure on traits that are less important in the domesticated context, such as camouflage and predator avoidance. Here, we hypothesize that strongly deleterious mutations may have become less deleterious in domesticated populations living alongside humans, while neutral mutations underlying traits of interest may have been selected by breeders, shifting the selection coefficients of genetic variants associated with these traits.

We test this hypothesis by studying the distribution of fitness effects (DFE) for nonsynonymous mutations in populations of wild wolves and breed dogs. The DFE is defined as the distribution of selection coefficients in an organism’s genome (or part of its genome) and it quantifies the proportions of new mutations that are neutral, deleterious, or beneficial (14). The DFE plays a fundamental role in population genetics, with implications for the understanding of the genetic architecture of complex traits, the evolution of recombination, and the survival of threatened species of ecological concern (14–17). Moreover, it also informs about the adaptive potential of a species and the amount of background selection expected in an organism’s genome (18–20). Despite its significance, our understanding of the biological factors influencing the DFE remains incomplete (21). Comparing different model organisms, Huber et al. (22) detected significant differences in the DFE across large evolutionary time scales, in particular between humans and flies. On a smaller time-scale, however, the DFE seems to be more stable. A comparative study on humans, flies, and tomatoes found that the DFE of different populations of the same species (or subspecies of the same genus) is highly correlated, with a correlation coefficient ranging from 0.91 to 0.99 (23). Importantly, their work found that the correlation is inversely related to genetic differentiation between populations. Similarly, Castellano et al. (24) found that the shape of the deleterious DFE is strikingly similar across great apes. Among the two main factors shaping the DFE are the organism complexity and the effective population size (*N_e_*) (22, 24–26). Organism complexity – defined in terms of a larger number of unique cell types coupled with a larger genome with more genes and more protein-protein interactions – should be similar in phylogenetically related organisms. Thus, all else being equal, we would expect the DFE of closely related species, like the ones examined in refs. (23, 24) to indeed be highly correlated and similar. However, the other piece of the puzzle, namely the *N_e_*, is known to vary across closely related species and even across populations within a species (24, 27) and thus, in principle, we can expect to see changes in DFE across short evolutionary time scales.

While our understanding of the determinants of the DFE is increasing, there are still many gaps such as how much it varies across non-model species and what factors in addition to organism complexity and *N_e_* shape the DFE. Insights from comparing the ratio of nucleotide diversity for nonsynonymous and synonymous variants (π_N_/π_S_) across plant species revealed additional factors that may influence the DFE (21, 28). These features include selfing/outcrossing, longevity, reproductive system, and ploidy ((21) and references therein). While the full DFE cannot be inferred from π_N_/π_S_ alone, as this ratio also is affected by changes in population size, these observations suggest that additional factors related to life-history traits could play a role in determining the DFE of a species.

Domestication can impact both the life-history traits and the *N_e_* of a species (29, 30), raising the question of whether it might also alter the DFE. Additionally, although the process of domestication can follow different pathways (31) and vary across and within species (1), it typically entails fast and drastic changes in the selective pressure acting upon a population (1, 7, 32). For instance, at the early stages of domestication, selection for tameness and against aggressiveness may take place as animals start living in the surroundings of human populations (31, 32). Shifts in dietary intake are also expected, since animals may take advantage of the resources associated with human societies such as food waste, smaller animals that are attracted to it, and surplus food (31, 33). At later stages of domestication, in particular, during breed formation and the improvement of traits, we expect to see fast shifts in the selective pressure (1). For instance, many of the traits of interest for breeders may be deleterious in the wild – such as an increase in tameness, extreme morphological alterations (e.g., brachycephaly), and the loss of camouflage – which means that, in practice, humans could be selecting for traits that otherwise would be eliminated in the wild. These phenomena combined, in particular the changes in *N_e_* and in the selective pressure resulting from intense artificial selection, could in principle result in a shift in selection coefficients, changing the DFE in domesticates relative to their wild counterpart. In this context, domesticated animals offer a unique opportunity for the study of the determinants of the DFE, specifically by allowing for dissecting the relative importance of drastic environmental shifts (emulated by the process of domestication and breed formation) and the shared organismal characteristics of closely related species (the wild and the domestic counterparts).

To tackle these questions, we focused on comparing the DFE of wild wolves and domestic dogs. Evidence suggests that dogs were domesticated from gray wolves (*Canis lupus*) around 15,000 years ago or earlier (29, 34, 35), making them the oldest known domesticated species. Modern dog breeds arose much more recently, around 200 years ago through multiple processes involving intense artificial selection, inbreeding, and gene flow (36). Both the initial domestication of dogs and the more recent development of dog breeds entailed severe population bottlenecks (29, 35–37), with significant consequences to the dog genetic diversity such as an excess of deleterious genetic variants (5) and runs of homozygosity (8). Here, we ask whether domestication has also shifted the selection coefficients in the dog genome relative to that of the wolf. We address this question by leveraging whole-genome sequence data from 24 gray wolves and 61 dogs. After accounting for differences in demography and background selection, we find that the DFE is strikingly similar across canids, with 26-28% of new amino acid-changing mutations being neutral/nearly neutral (|*s*| < 1e-5), and 41-48% under strong purifying selection (|*s*| > 0.01). We evaluate the robustness of our results to different model assumptions and conclude that the DFE is stable across short evolutionary timescales, even in the face of putative drastic changes in the selective pressure caused by artificial selection during domestication and breed formation. On par with previous works describing DFE evolution across the tree of life, our results indicate that the DFE of nonsynonymous mutations depends more strongly on genome structure and organismal characteristics, and less so on shifting selective pressures or environmental factors.

## Results

### Genetic Diversity in Wolves and Breed Dogs

We analyzed four canid populations for which publicly available, high-coverage (>30x), whole genome sequences were available: Arctic Wolf (AW; n = 15 (38, 39)), Border Collie (BC; n = 10 (40)), Labrador Retriever (LB; n = 10 (40)), and Pug (PG; n = 15 (41)). Genomic VCF files were generated for each population according to the pipeline outlined by Phung et al. (39) (https://github.com/tanyaphung/NGS_pipeline). These VCFs were subset to exonic regions considering the CanFam3.1 reference genome exon annotations. We exclusively considered biallelic SNVs where all three potential variants were annotated as either synonymous or missense (henceforth nonsynonymous) mutations, in practice excluding sites with potential nonsense mutations, splice sites variants, and indels. This resulted in a total exonic sequence length of ∼21.7 Mb for which we retrieved data in the four canid populations.

Genetic variation data was summarized by the folded site frequency spectrum (SFS) for each population, after the removal of related individuals and projecting down the sample size in order to maximize the number of usable SNPs. No evidence of substantial population structure within each group was detected based on a Principal Component Analysis (Fig. S1). The final folded SFSs are shown in Fig. S2 and S3, highlighting nonsynonymous variants segregating at lower frequency relative to synonymous mutations.

### Controlling for the Effects of Demography and Background Selection in the Estimation of the DFE

In this work, we sought to characterize the effects of changing selective pressures (caused by artificial selection during the process of domestication and breed formation) on the DFE of new mutations. To do so, we initially used the SFS of the wolf (AW) and three breed dog populations (BC, LB, and PG) mentioned above. The SFS of a population is shaped by different evolutionary forces such as genetic drift, natural selection, and background selection in linked sites. To untangle the effects of target natural selection from the effects of demographic changes and background selection, we followed an approach developed by Kim et al. (42) consisting of two steps. First, we use the SFS of synonymous variants to infer the underlying demography; and second, we use the inferred demographic model and the SFS of nonsynonymous variants to infer the DFE. We implemented this approach with *∂a∂i* (43), a maximum-likelihood method that uses diffusion approximations to fit population genetic models of demographic history and natural selection to genetic polymorphism data summarized in SFS.

We initially considered a two-stage demographic model (henceforth, “2-epoch”) allowing for one instantaneous population size change, and used the multinomial likelihood to infer the demographic parameters. The parameters of the model are *ω* – the intensity (i.e., fold-change) of the population size change – and time *T* that this demographic change occurs in the past. The method also outputs *θ_S_*, the estimated population mutation rate for synonymous mutations, defined as *θ_S_*=4\**N_a_*\**μ*\**L*, where *N_a_* is the estimate of the effective population size of the ancestral population (before the population size change), *μ* is the mutation rate per site, per generation, and *L* is the coding sequence length. The inferred demographic model parameters are presented in Table S1. We infer the wolf’s population *N_a_* at ∼72,000 individuals and a population size reduction (*ω*) of approximately 19% of its size at ∼2,000 generations ago. Estimates for BC and LB are in the same order of magnitude, with *N_a_* = ∼52,000 and ∼86,000, *ω* = ∼34% and ∼20% respectively, although population size change takes place at an earlier period (∼17,500 generations ago in either population). The estimates for PG indicate a much larger ancestral population effective size of ∼223,000 and a much more severe bottleneck (*ω* = 3.5%), taking place more recently (∼8,000 generations ago). Both the model and the data SFSs, as well as the residuals of the model fit, are shown in Figures S4 and S5, highlighting the models fit the data well.

**Stability of the DFE Despite Domestication.** We next inferred the DFE for new mutations using the nonsynonymous SFS, conditioning on the maximum-likelihood demographic parameters inferred from the synonymous variants. In doing so, we do not directly quantify the selection coefficient of each variant, but instead, summarize the distribution of fitness effects (DFE) over many sites. Initially, we focused on all exons annotated in the CanFam3.1 reference genome assembly for this inference, in order to calculate the genome-wide DFE of nonsynonymous mutations for each population of canids.

To infer the DFE of nonsynonymous mutations, we used *fit∂a∂i* (42), a modification of *∂a∂i* (43) that allows for the inference of DFE from polymorphism data. Initially, we considered that the selection coefficients (*s*) followed a gamma distribution and inferred its shape and scale parameters. In addition to the gamma distribution, at a second stage, we also considered a mixture distribution where a proportion of mutations are neutral (*s* = 0) and the rest follow a gamma distribution (“neugamma” henceforth). We effectively treat the neugamma distribution as a single integrable function, as described in ref. (42).

The inferred gamma-distributed DFE in canids (Fig. 1) shows no significant differences across the wolves (AW; yellow) and three different breeds of dogs (BC, LB, and PG; different shades of blue). The proportion of neutral/nearly-neutral mutations (|*s*| < 1e-5) is ∼26-28% across wolves and dogs and that of strongly deleterious mutations (|*s*| > 1e-2) is ∼41-48% (Table S2). No statistically significant differences between the discretized DFE were observed across canid populations, based on the overlap of the 95% confidence intervals (Fig. 1).

**Fig. 1.**
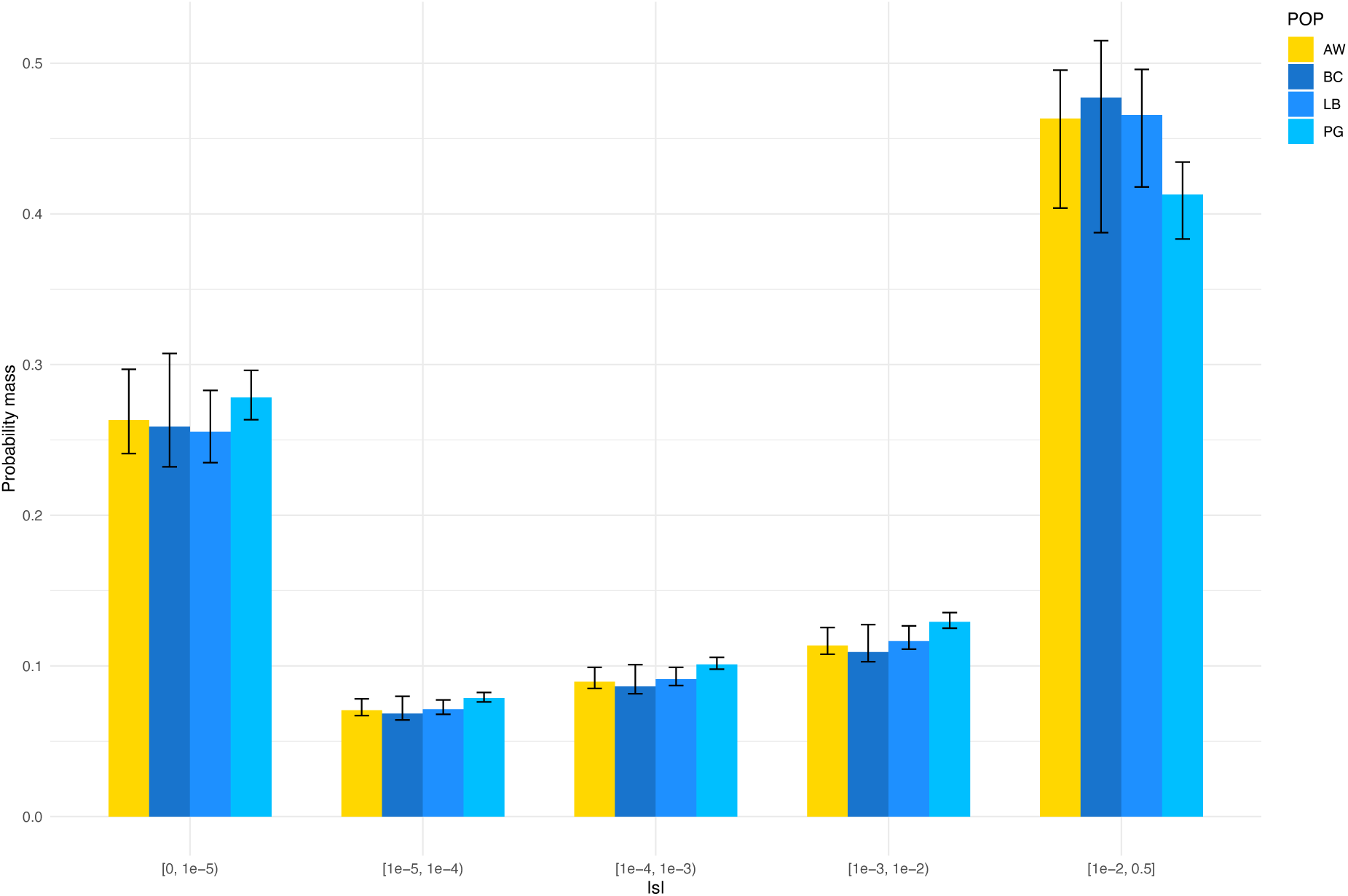
Discretized distribution of fitness effects (DFE) showing the proportions of nonsynonymous mutations in various categories of |*s*|. From left to right, mutations range from neutral/nearly neutral (0 < |*s*| ≤ 1e-5) to strongly deleterious/lethal (|*s*| ≥ 1e-2). The DFE is assumed to follow a gamma distribution. The arctic wolf population (AW) is depicted in yellow, and three domestic dog breeds in different shades of blue (BC = border collie; LB = labrador retriever; PG = pug). Error bars represent the 95% confidence intervals for the proportion of mutations in each category of |*s*|.

Assuming a neugamma distribution for the DFEs, we observe a similar pattern, although the confidence intervals are considerably larger (Fig. S6). While the neugamma distribution improves the fit of the model for some populations (Table S3), we found that the optimizations for the neugamma did not converge as well as the ones for the former. On a related note, when we compare the DFEs of these four canid populations considering the distribution with best log-likelihood in each case (namely gamma for BC and neugamma for the other three populations), we still see no significant differences in the DFEs across populations when considering the 95% confidence interval (Fig. S7), although the neugamma DFE for PG predicts a considerably higher proportion of strongly deleterious mutations than that in the other populations. The estimated parameters of the gamma and neugamma DFEs for the analyses, as well as the corresponding log-likelihoods, are reported in Tables S2 and S3. The model SFS fits for the nonsynonymous SFS for both the gamma and neugamma distributions and corresponding residuals are shown in Figures S8-S11, indicating the inferred DFE models fit the data well considering either functional form of the DFE.

We observed a similar pattern when we use the same sample sizes of n_eq_ = 6 across all four populations (Fig. S12), although the 95% confidence intervals are wider, as expected due to the reduced data. Finally, we computed the population-scaled DFE, *2N_a_s*, for the four canid populations. Estimates of *2N_a_s* measure the relative strengths of selection vs. drift acting in the population and are more robust than estimates of *s*, as *2N_a_s* is directly inferred from *fit∂a∂i*. Similar to the population-scaled DFE in terms of *s*, we detect no significant differences in the DFE across AW, BC, LB, and PG (Fig. S13). In sum, the inference of a stable DFE for nonsynonymous mutations in canids is not an artifact caused by different sample sizes across populations or biases in converting estimates of *2N_a_s* into estimates of *s*.

### The Wolf and Dog DFEs are Highly Correlated

Because dogs and wolves have diverged relatively recently (29, 34, 35), and we did not observe any differences in the DFE between dogs and wolves, we sought to jointly model their DFE, following the approach developed by Huang et al. (23). According to this approach, the joint DFE is estimated for pairs of populations, and the correlation of their selection coefficients (*ρ*) is estimated. That is to say, *ρ* = 1 means that the mutations have the same selection coefficients in both populations, while lower values of *ρ* indicate that those mutations that are deleterious in one population may not be deleterious in the second population. This approach follows the same framework of *∂a∂i/fit∂a∂i*, using the synonymous SFSs to control for the effects of demography and background selection, and the nonsynonymous SFSs for the DFE inference, except that it uses the joint SFS (i.e., 2-dimensional; henceforth “2D”), computed for a pair of populations (23). The method requires only modest sample sizes and is robust to many forms of model misspecification (23).

We first fit a simple demographic model (“split_mig”) with a population split at time *T* in the past (Fig. S14-S16). According to this model, the size of the two derived populations relative to the ancestor is *ω_1_* and *ω_2_*, and remain constant after the split. Symmetric migration between the two derived populations happens at a rate *m*. We consider the following pairs of populations: AW-BC, AW-LB, and AW-PG. Projected sample sizes are the same used for the single population analysis described above. The joint SFS and fit plots are found in Figures S14-S16 and the inferred demographic parameters, in Table S4. The 2D demographic models show similar joint inferred demography across each pair, with *N_a_* = ∼20,500, effective population size in the present (*N_p_*) for the wolf population after the split at ∼8,400, and split time at ∼4,000 generations ago. The only noticeable difference across the three models is the *N_p_* for the breed dog populations after the split, which ranges from ∼500 for PG to approximately 1,300 for BC and LB (Table S4).

We inferred the joint DFE using the joint SFS of nonsynonymous variants (Fig. S17-19). Since little is known about the joint DFEs of wolf and breed dog populations, we first considered a simple bivariate lognormal distribution with an easily interpretable correlation coefficient. Given the similarity in DFEs for single populations, we assumed a symmetric bivariate lognormal distribution in which the marginal DFE for the wolf and dog populations are the same, while the parameter *ρ* quantifies the correlation of fitness effects of mutations between populations. Using a Poisson likelihood framework, we inferred the DFE parameters for each wolf-dog population pair (Table S4). We found that the DFEs of wolf and dog across all three comparisons are perfectly correlated (*ρ* > 0.99; Table 1). The discretized DFE for the bivariate lognormal joint DFEs and single-population gamma DFEs predict similar proportions of mutations in each bin (Table 1), suggesting the robustness of our findings relative to the assumptions about the functional forms of the DFE and whether we model the populations separately or jointly.

**Table 1.** Comparison of the proportions of nonsynonymous mutations in various categories of |*s*|. Shown are the estimates for the Arctic wolf (AW), border collie (BC), labrador retriever (LB), and pug (PG) considering the 1D site frequency spectrum (SFS) and for each pair of populations including AW and each of the three dog breeds based on the 2D SFS. The AW’s DFE is assumed to be gamma-distributed, while the joint DFEs are assumed to be lognormal distributed. ^a^*ρ* represents the correlation coefficient of the DFE between pairs of populations. ^b^F_ST_ from ref. (27).

| Population comparison | Proportions of nonsynonymous mutations in various categories of $\vert s\vert $ | | | | | $\rho^a$ | $F_{ST}^b$ |
| --- | --- | --- | --- | --- | --- | --- | --- |
|  | [0, 1e-5) | [1e-5, 1e-4) | [1e-4, 1e-3) | [1e-3, 1e-2) | [1e-2, 0.5) |  |  |
| AW | 26% | 7% | 9% | 11% | 46% | n/a | n/a |
| BC | 26% | 7% | 9% | 11% | 48% | n/a | n/a |
| LB | 26% | 7% | 9% | 12% | 47% | n/a | n/a |
| PG | 28% | 8% | 10% | 13% | 41% | n/a | n/a |
| AW-BC | 20% | 8% | 9% | 10% | 53% | 0.999 | 0.166 |
| AW-LB | 21% | 7% | 8% | 8% | 56% | 0.999 | 0.175 |
| AW-PG | 21% | 8% | 9% | 9% | 54% | 0.999 | 0.234 |

### Assessing the Robustness of the Inferred DFE

In order to confirm our observations about the stability of the DFE despite domestication, we sought to replicate our results using a separate dataset comprising one wolf and two dog populations. The wolf population (MW) comprises the genomes of nine gray wolves sequenced at ∼19x depth of coverage (5). The two dog populations comprise one sample with 20 domestic breed dogs (MD) sequenced at an average of ∼18x (5) and 10 Tibetan mastiffs (TM) sequenced at ∼15x (39). Only one individual was removed from TM due to relatedness with other TMs in the sample. Following the same approach used for the higher coverage data (namely AW, BC, LB, and PG), we projected down the sample size in order to maximize the number of usable sites. The final sample sizes after projection and removal of related individuals are MW = 8, MD = 16, and TM = 7.

We fit a 2-epoch demographic model to the synonymous SFS (Table S1; Fig. S4 and S5) and a gamma-distributed DFE model to the nonsynonymous SFS (Table S2; Fig. S8 and S9) using*∂a∂i/fit∂a∂i*. Although the maximum likelihood estimates for the proportion of new mutations with different values of *s* may appear to differ between wolves (MW) and dogs (MD and TM) using this lower coverage dataset (Fig. S20), the 95% confidence intervals of these estimates overlap, suggesting no significant differences between the DFEs of MW, MD, and TM. This observation confirms our findings obtained with the analysis of the high coverage samples (Fig. 1) showing no significant differences in the DFE of wolves and dogs.

In addition to confirming the observations of a stable DFE in canids with an independent dataset, we also assessed whether misspecification in the model parameters could bias our estimates. We did so by using the high coverage samples (AW, BC, LB, and PG), considering their full, projected sample sizes, and re-ran the analyses considering a low mutation rate of 3.00e-9 (considering the lower range of the estimate from ref. (35)), a high mutation rate of 6.73e-9 (considering the cat-dog divergence-based mutation rate from ref. (39)) and a 1.25x higher mutation rate in exons – see Materials and Methods), the expected ratio of nonsynonymous-to-synonymous mutations (NS:S) estimated for humans (42), and a 3-epoch demographic scenario (Table S5).

The inferred DFE for dogs and wolves in each case are highly similar (Fig. S21), confirming our observations are robust to misspecification of the model, as far as it concerns the mutation rate (within a reasonable range for canids (35, 39)), the expected NS:S ratio (within a reasonable range for mammals (42)), and the number of population size changes. We note that the log-likelihood of the inferred 3-epoch demographic model for PG is slightly improved relative to that of the 2-epoch model (log-likelihood = -79.59 and -72.95 respectively); however, the gamma DFE for PG is qualitatively similar regardless of the demographic model considered (Fig. S22).

### DFE Inference for Domestication-associated Genes

While our results using the whole dog exome point to no differences in the DFE of nonsynonymous mutations in dogs and wolves (Fig. 1), even when considering different model assumptions and population samples (Table 1, Fig. S6, S12, S20, and S21), we hypothesized that the effects of domestication on the DFE may be more pronounced in gene sets thought to be associated with domestication. Although no single suite of traits is consistently seen across domestic animals (44), the literature harbors a number of studies evidencing signals of natural selection in domesticated species associated, in particular, with the nervous system, behavioral traits, skeletal system development, immunity, and pigmentation (7, 12, 13, 29, 32, 45, 46). Thus, to tackle this question, we subset the whole-exome data to different sets of genes implicated in pathways putatively associated with domestication.

To implement this approach, the whole-exome data were filtered based on the following Gene Ontology terms: Nervous System Development (5.5 Mb exonic sequence length), Immune System Processes (3.6 Mb), and a combination of Immune System Processes, Nervous System Development, Carbohydrate Metabolic Processes, Pigmentation, and Skeletal System Development (“Domestication Genes” subset; 10.4 Mb). Similarly to our previous analyses, we fit first the synonymous SFS from the genes in each set in AW, BC, LB, and PG (considering the maximum sample size projected) to 2- and 3-epoch demographic models. Based on log-likelihoods estimated by *∂a∂i*, we picked the best demographic model for each population considering each gene set independently for a total of 12 models (Tables S1 and S5). We assume a gamma-distributed DFE and fit the resulting model SFS to the data nonsynonymous SFS following the same procedure adopted for the whole exome analysis.

The discretized gamma-DFE computed for each gene set shows no significant differences in the predicted proportion of deleterious and neutral mutations at different selection strengths (Fig. 2). While small differences in the maximum-likelihood estimates for these proportions are visually evident in some cases, in particular showing a larger proportion (additional ∼10%) of strongly deleterious mutations in PG relative to the others for the Nervous System Development and “Domestication Genes” sets, the 95% confidence intervals largely overlap (Table S6).

**Fig. 2.**
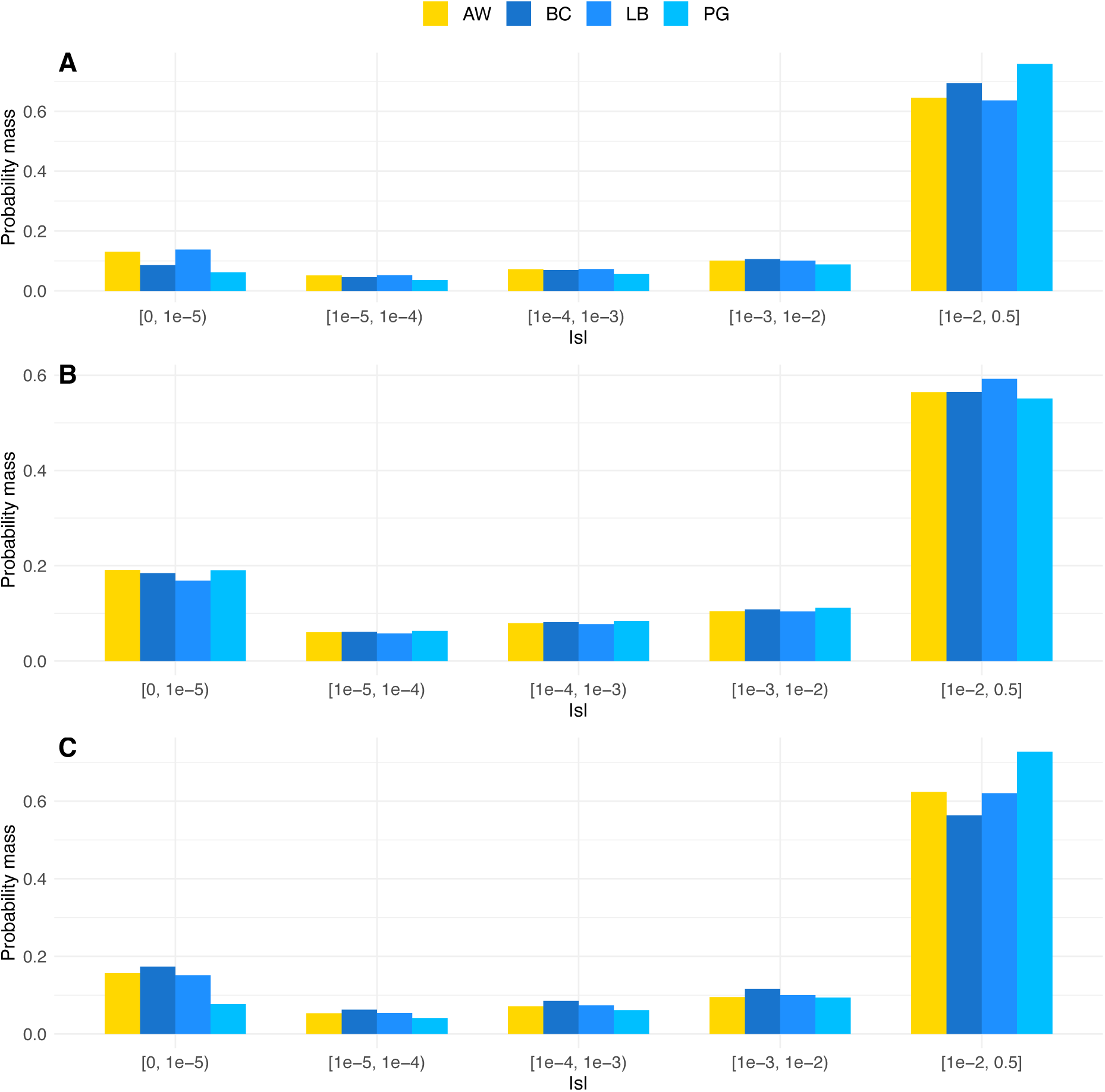
Discretized distribution of fitness effects (DFE) showing the proportions of nonsynonymous mutations in various categories of |*s*| for different gene sets: (A) Nervous System Development, (B) Immune System Processes, and (C) a combination of Immune System Processes, Nervous System Development, Carbohydrate Metabolic Processes, Pigmentation, and Skeletal System Development (“Domestication Genes” subset). In all panels, mutations range from neutral/nearly neutral (1e-5 < |*s*| ≤ 0) to strongly deleterious/lethal (|*s*| ≥ 1e-2). The DFE is assumed to follow a gamma distribution. The arctic wolf population (AW) is depicted in yellow, and three different domestic dog breeds in different shades of blue (BC = border collie; LB = labrador retriever; PG = pug). Confidence intervals for these estimates can be found in Table S6.

## Discussion

### The Evolutionarily Stable DFE of Dogs and Wolves

Our findings allow us to revisit our initial hypothesis that shifts in selective pressures during domestication would change the DFE in domesticated species compared to their wild relatives. We specifically hypothesized that strongly deleterious mutations might have become less deleterious under domestication and that this would have an impact in the DFE of new amino acid changing (nonsynonymous) mutations. At the early stages of domestication, selection for tameness may take place as animals start living in the surroundings of human populations (31, 32). At later stages, we expect to see fast shifts in the selective pressure (1), with many of the traits selected by breeders being potentially deleterious in the wild. Comparing populations of wolves and domestic dogs, we sought to test whether these putative shifts in selective pressure would also have an impact on the DFE of dogs. Contrary to our initial expectations, our data showed striking stability of the DFE across wild and domestic canid populations (Fig. 1), suggesting that domestication and the development of breeds may not drastically alter selection coefficients in nonsynonymous mutations, as inferred from polymorphism data. We assessed the robustness of our findings relative to differences in sample sizes across populations (Fig. S12) and model assumptions (Fig. S6 and S21; Table 1), and replicated our findings with an independent dataset (Fig. S20), and consistently observe statistically similar DFEs across wolves and breed dogs.

This stability in the DFE across wolves and dogs aligns with previous studies comparing DFE evolution across the tree of life, which have found that significant differences in the DFE occur primarily across distantly related species (22, 47), whereas minimal changes are observed among more closely related species, subspecies, or populations (23, 24, 48, 49). Insights from comparing the ratio of nucleotide diversity for nonsynonymous and synonymous variants (π_N_/π_S_) across plant species also pointed to a limited role of domestication on shifting selection coefficients in plants (21, 28). The observed stability of the DFE within short evolutionary time scales supports the idea that the DFE is influenced more strongly by intrinsic organismal characteristics (e.g., organismal complexity, genome structure, genetic context, life history, etc.) than by extrinsic environmental factors (21, 22, 28), here, emulated by domestication.

Despite evidence showing stability of the DFE, in particular in closely related species and populations (23, 24, 48, 49), experimental evolution studies in *Drosophila* (50) and the bacterium *E. coli* (51) indicate that environmental shifts can influence the DFE. In particular, Wang et al. (50) showed that especially the variance, V(*s*), of the DFE of *Drosophila* was dependent on the environment. On a similar note, the strength of natural selection on pigmentation genes was found to be different across human populations (52). Thus, while the effects of domestication and breed formation on the DFE of canids appear minimal in our study, environmental factors may still play a role under certain evolutionary contexts. Motivated by these observations, we sought to investigate the effects of domestication on gene sets in biological pathways thought to be impacted by domestication in animals, in particular, the nervous system, skeletal system development, immunity, metabolism/diet, and pigmentation (7, 12, 13, 29, 32, 45, 46). Surprisingly, we found that the DFE is stable even when analyzed for these gene sets (Fig. 2), suggesting artificial selection may not have affect the DFE of canids, even for domestication-associated gene sets.

One potential explanation for the observed DFE stability is that domestication-related shifts in selective pressure have not been sufficient to generate detectable changes in the DFE, at least within the sample sizes and timescales studied. Forward simulations by Castellano et al. (53), modeling pig domestication over 10,000 generations, suggest that the DFE of deleterious mutations can be accurately estimated using either the 1D and 2D SFSs–that is either modeling one population at a time or two populations jointly. In their simulations, the evolutionary effects produced by domestication are modeled by changing, at the time of the split, the fitness effects of a proportion (5% or 25%) of the existing and new mutations in the domestic population relative to the wild counterpart. Given their findings, our research design appears to be well-powered for detecting meaningful DFE changes in canids, assuming similar dynamics to what they simulated also apply to dog domestication.

Alternatively, domestication might have more pronounced effects on other mutation types, such as regulatory variants or complex structural variants, rather than the nonsynonymous mutations studied here. Indeed, variants that differentiate wolf and domestic dog populations are enriched in regulatory elements such as promoters and enhancers (54), suggesting that a promising area for further investigation is to look at DFE differences between dogs and wolves focusing on regulatory variation. Finally, it is possible that domestication and artificial selection during dog breed formation have had a greater impact on beneficial mutations (55) or standing deleterious variation, which we did not specifically assess in this study. Further analyses targeting beneficial DFE components, as well as standing variation, may provide additional insights into the evolutionary consequences of domestication in canids.

We note that the sample sizes used for our inference, ranging from *n* = 6 (*n_eq_*) to 16 (MD after projection), may also limit power to detect subtle shifts in natural selection, as strongly deleterious variants are expected to segregate in low frequencies and thus not be observed in our sample. Instead, the strongly deleterious part of the DFE is extrapolated from the lack of common variants and the functional form of the DFE assumed. Larger sample sizes including more rare variants could enable more accurate inference of the more strongly deleterious part of the DFE. However, we note that estimates of the proportion of neutral/nearly neutral variants (|*s*| < 1e-5) among new mutations is less likely to be impacted by sample size. For this portion of the DFE, where we have more information, we do not detect a difference across wolves and breed dogs. This finding suggests that if we assume the DFE of new mutations is gamma distributed, we can use data from a small number of individuals to learn about mutations that we did not observe in our sample.

### The canid DFE relative to other animals

A recent study analyzed the DFE across eleven animal (sub)species, including humans, mice, fin whales, vaquitas, gray and Arctic wolves, collared flycatchers, pied flycatchers, halictid bees, *Drosophila*, and mosquitoes (56). Their findings show variation in the DFE across deep evolutionary time, with mammals having a larger proportion of strongly deleterious (|*s|* > 1e-2) mutations (22% in vaquitas to 47% in Arctic wolves) than other animals (0.0% in *Drosophila* to 5.4% in collared flycatchers), while the proportion of weakly deleterious mutations (1e-5 ≤ |*s|* < 1e-3) is smaller in mammals relative to birds and insects (56). Previously, Huber et al. (22) examined five competing models explaining the determining factors driving DFE evolution and found strong support for the Fisher’s Geometrical Model (FGM). According to the FGM, phenotypes are characterized as points in an *n*-dimensional space, with fitness being a decreasing function of the distance from the optimal phenotype (57). The phenotype dimensionality *n* can be understood as the “organismal complexity.” Among others, the FGM makes one key prediction that is confirmed by refs. (22, 56) that mutations in more complex organisms are on average more deleterious because they are more likely to disrupt an important function in a complex organism than in a simpler one. In this context, the unusually high proportion of strongly deleterious mutations (|*s|* > 1e-2) in canids inferred in this study (41-48%) relative to other mammals (30% fin whale, 27% human, 24% mouse, and 22% vaquita (56)) suggest a venue worthy of further investigation.

### DFE and Homologous Recombination in Canids

We speculate that our findings on the large amount of predicted strongly deleterious mutations in dogs and wolves relative to other mammals, including humans, may relate to the unique recombination landscape in canids, where recombination predominantly occurs in promoter regions due to a non-functional *PRDM9* gene (58). This recombination pattern differs significantly from other mammals, where the PRDM9 protein localizes recombination hotspots throughout the genome (59, 60). In species harboring a functional copy of *PRDM9*, recombination hotspots may localize anywhere in the genome. The restriction of recombination to promoter regions in canids may thus effectively reduce recombination within genes, potentially impacting the purging of deleterious mutations under Muller’s Ratchet hypothesis, which posits that low-recombination regions are more prone to accumulating harmful mutations (61, 62). We note that less efficiently purging deleterious mutations does not necessarily equate to a more deleterious DFE. However, because our inference is based on polymorphism data, linkage between mutations within genes could skew the SFS, resulting in a larger proportion of deleterious mutations being inferred.

As a first step into examining the relationship between the DFE and recombination rates, we inferred an LD-based recombination map for the Arctic wolf and subset its genome into three different datasets based on the estimated recombination rates (*r*), in units of recombination events per bp per generation: Low (0 ≤ *r* < 1.9e-9), Moderate (1.9e-9 ≤ *r* < 4.3e-9), and High (*r* ≥ 4.3e-9) recombination rate. We then used these three subsets to infer the DFE using the same framework implemented for the whole exome. Our results show qualitatively similar DFEs between low and high recombination regions, with regions with moderate recombination rate presenting an overall larger proportion of deleterious mutations (Fig. S23). One potential explanation for this observation is that intermediate recombination rate regions could have genes with different functions than other portions of the genome. We note that, due to the small sample sizes considered in our study, our findings regarding the differences in the proportion of strongly deleterious mutations across regions with different recombination rates should be considered with care. On the other hand, estimating the proportions of nearly neutral variants among new mutations is less likely to be impacted by sample size, and in the case of nearly neutral sites (*|s*| < 1e-4), we inferred highly similar proportions for the whole exome (33%) and the different recombination subsets (Low *r* = 34%; Moderate *r* = 29%; High *r* = 32%). Given the importance of recombination in shaping genetic variation, understanding how the unique canid recombination pattern within mammals affects the DFE could provide valuable insights into the determinants of the DFE and the implications of PRDM9-independent recombination on adaptive potential and the purging of detrimental variants.

## Conclusion

Overall, our study provides evidence that the DFE of nonsynonymous mutations remains relatively stable despite the significant shifts in selective pressures putatively associated with domestication in canids. Our finding underscores the role of intrinsic organismal characteristics in shaping the DFE, while environmental factors and shifts in the selective pressures associated with domestication may have limited influence – contradicting previous findings in *Drosophila* and *E. coli* showing an influence of the environment in shaping the selecting effects of nonsynonymous mutations (50, 51). Future studies could explore the full properties of the DFE in other domesticated species with varying degrees of selection intensity and different life history traits to determine whether these observations hold across domesticated lineages. Additionally, investigating the DFE of gene promoters and enhancers in domesticated species could yield further insights into the importance of natural selection on regulatory variation during domestication. In conclusion, while domestication has clearly impacted genetic diversity and allele frequencies in canids (and other domesticated species alike), our data suggest that the DFE of nonsynonymous mutations remains resilient to these shifts, providing new perspectives on the stability of fitness effects even in the face of drastic environmental shifts.

## Materials and Methods

### Genomic Data

Whole-genome sequencing data were aggregated from the literature: Arctic wolf (AW; n = 15) with ∼39x coverage (38, 39), border collie (BC; n = 10) with ∼24x coverage (40), labrador retriever (LB; n = 10) with ∼30x coverage (40), pug (PG; n = 15) with ∼47x coverage (41), and Tibetan Mastiff (TM; n = 10) with ∼15x coverage (39). In addition to these, we also aggregated data from nine wolves from different populations (referred to as “mixed wolves” or MW in this study) with ∼19x coverage and 20 dogs from 20 different breeds (referred to as “mixed dogs” or MD in this study) with ∼18x coverage (5). The term “mixed” here refers to the fact that these individuals came from different populations and were pooled into one sample, not that they are necessarily wolf-domestic dog hybrids or mixed breed dogs.

Raw whole genome sequences (*fastq* files) were processed following GATK best practices, and according to the pipeline outlined by Phung et al. (39) (https://github.com/tanyaphung/NGS_pipeline). In brief, the *fastq* files were first aligned to the dog genome (CanFam3.1) with *BWA* (63). We then marked duplicate reads with Picard tools (https://broadinstitute.github.io/picard/), removed reads with mapping quality (MAPQ) less than 30 using *SAMtools* (64), and recalibrated the base quality scores using the *BQSR* tool in *GaTK* v3.8 (65, 66). We performed joint genotyping with the *HaplotypeCaller* method and emitted all sites (variant and invariant). To reduce bias in SNP calling accuracy between canids from a given dataset, we conducted joint genotyping considering all individuals in each population. For example, joint genotyping was conducted on the 15 AW samples as a group, separately on the 10 BC samples as a group, and so on. We then applied post hoc filtering to each of the VCF files generated for each of the seven datasets. Specifically, we applied GATK filtering recommendations for variant sites in non-model species: QD < 2.0, FS > 60.0, MQ < 40.0, MQRankSum < -12.5, ReadPosRankSum < -8.0, and a minimum genotype quality (GQ) of 20. Additionally, we removed clustered SNPs (i.e., > 3 SNPs within 10 bp).

For invariant sites, for which no best practices were available, we applied the following filters: QUAL < 30, and RGQ < 1. For both variant and invariant sites, we applied a minimum depth filter of 10 for each genotype, as previous work has found heterozygous calls are unreliable below this depth (5), and a maximum depth filter of 2.5 times the average genomic coverage (specific for each of the seven datasets). Finally, we removed any sites where all individuals were heterozygous, fewer than 80% of individuals in a group had a genotype call after post hoc filtering was applied, or any sites within the UCSC repeat regions.

Genomic VCF files were subset to autosomal exonic regions with *VCFtools* (67). Exon coordinates for the reference dog genome *canFam3* were obtained from http://hgdownload.soe.ucsc.edu/goldenPath/canFam3/database/ensGene.txt.gz (accessed on 02/05/2018). From this dataset, we obtained the coordinates for the exons of 30,784 genes, considering the longest transcript in each case. Based on the *canFam3* reference genome, we calculated the total exon length for dogs as 25.16 Mb.

To annotate the effects of all potential exonic single nucleotide variants (SNVs), we artificially introduced “mutations” to a VCF file containing dog exome data, so that all three potential SNVs were observed in each site. The functional effects of each variant in each site were predicted with *SnpEff* (68, 69), using the dog reference genome build CanFam3.1.75, available with *SnpEff*. We used these annotations to classify each exonic position into either a 0-, 2-, 3-, or 4-fold degenerate site. We exclusively considered exonic sites where all three potential SNVs were annotated as either synonymous or nonsynonymous (missense) mutations (∼21.7 Mb), effectively discarding sites with other types of annotations, such as splice sites and nonsense mutations (i.e., stop-gained and stop-lost).

### Computing the Site Frequency Spectra

We used *KING* (70), implemented in *PLINK* (71), to identify pairs of related individuals and excluded those with more than 35% of their genome with at least one allele in IBD in practice removing first-degree relatives (i.e., parent-child and siblings). Relatedness was estimated using a set of putative neutral SNVs from ref. (39). To select putative neutral regions in the dog genome, these authors filtered out any locus 0.4 cM away from conserved regions, annotated with *phastConsElements100way* UCSC Genome Browser, or a gene, resulting in approximately 24.5 Mb of sequence. The following individuals were excluded from the downstream analysis after removing related individuals: AW14, BC2, BC8, BC10, and TM4. The final sample sizes obtained for each population after the removal of one of the relatives were AW = 14, BC = 7, LB = 10, PG = 15, MW = 9, MD = 20, and TM = 9.

Because we allowed some missing data in our dataset and considering we cannot use missing data to calculate the SFS, we projected down these sample sizes in order to maximize the number of SNVs available for each population. We used *EasySFS* (https://github.com/isaacovercast/easySFS; (43)) to calculate the number of SNVs for a given sample size and to project down the sample size. *EasySFS* averages allele absolute frequencies over all possible combinations of samples for a given sample size (i.e., the hypergeometric projection method). The resulting projected sample sizes were: AW = 13, BC = 6, LB = 9, PG = 14, MW = 8, MD = 16, and TM = 7. Additionally, to assess if unequal sample sizes were biasing results, we further projected down these sample sizes to n_eq_ = 6 for the high coverage samples (AW, BC, LB, and PG).

Based on these sample sizes after filtering and projection, we calculated the folded SFS for each population using *EasySFS*. The folded SFS describes the number or proportion of variants at different minor allele frequencies in the sample. We chose to use the folded SFS (as opposed to the unfolded) to avoid biases resulting from the misspecification of the ancestral allele (72).

### Calculating Synonymous and Nonsynonymous Sequence Lengths

After classifying each exonic position into 0-, 2-, 3-, or 4-fold degenerate sites, we calculated the ratio of nonsynonymous sequence length to synonymous sequence length (L_NS:S_) in the *canFam3* genome assembly. We used L_NS:S_ to calculate the expected number of nonsynonymous mutations from the inferred synonymous mutation rate for the DFE inference (see below).

The nonsynonymous sequence length (L_NS_) was calculated as the number of 0-fold degenerate sites, ⅔ of the 2-fold degenerate sites, and ⅓ of the 3-fold degenerate sites. Conversely, the synonymous sequence length (L_S_) was calculated as the number of 4-fold degenerate sites, ⅔ of the 3-fold degenerate sites, and ⅓ of the 2-fold degenerate sites. Because methylated CpG sites are highly mutable (73, 74) and enriched in exons (as seen in humans (75)), we calculated L_NS:S_ considering a 10x higher mutation rate in putatively methylated CpG sites. We defined putatively methylated CpG sites as those CpG sites not in CpG islands, which are known to be unmethylated (76, 77). CpG islands coordinates in the dog genome were obtained from http://hgdownload.soe.ucsc.edu/goldenPath/canFam3/database/cpgIslandExt.txt.gz (accessed 03/20/2018). This information was used for exploring the effects of different L_NS:S_ in the estimates of the DFE in different populations, using L_NS:S_ = 2.21 calculated for the dog (this study) and L_NS:S_ = 2.31 calculated for the human genome (22).

### Demographic and DFE Inference

We inferred demography and the DFE from site frequency spectrum (SFS) according to a maximum likelihood approach in two steps. First, we inferred the parameters of a demographic model considering synonymous variants. Second, we inferred the parameters of the DFE of nonsynonymous variants conditional on the demographic inference. As shown by Kim et al. (42), this two-step approach effectively controls for the effects of demography and background selection when estimating the DFE of nonsynonymous variants. The rationale behind this approach is that, if nonsynonymous variants are completely neutral, the SFS computed based on these variants will have the same shape as the SFS of synonymous variants and the number of nonsynonymous mutations will be 2.21x larger (for dogs, as calculated in the present study). Any differences between the shapes of the synonymous and nonsynonymous SFSs or a deviation in the number of mutations from the expected can be attributed to the effects of natural selection. That is because both synonymous and nonsynonymous variants are subject to the same demography. Note that, although the demographic model inferred in the first step may be biased by linked selection (78), using this combination of synonymous and nonsynonymous variants allows us to control for the effects of background selection in addition to demography when inferring the DFE in the last step (22, 42).

We implemented both the demographic and DFE inferences using *varDFE* (https://github.com/meixilin/varDFE; (56)), a robust but flexible workflow implemented in Python. The demographic inference was performed with *∂a∂i* (43), implemented via *varDFE* (*Demog1D_sizechangeFIM* module). The method implemented via *∂a∂i* uses a diffusion approximation to compute the SFS given a demographic model. The multinomial likelihood is maximized to estimate the demographic parameters from the observed (data) synonymous SFS. A population mutation rate for synonymous variants *θ_S_* is estimated by scaling the optimized SFS relative to the observed synonymous SFS. The ancestral population size *N_a_* can then be estimated considering *θ_S_* according to this equation: *θ_S_*=4\**N_a_*\**μ*\**L. varDFE* uses *N_a_* for scaling the time and size parameters of the demographic model as well as the selection coefficients inferred at the final step.

We considered two simple demographic models with spontaneous size changes. The 2-epoch model has a single size change and the 3-epoch model has two size changes. As can be seen in Figures S4 and S5, the SFSs computed from these simple demographic models present a good fit to the observed synonymous SFSs.

We used *fit∂a∂i* (42), implemented via *varDFE* (*DFE1D_refspectra* and *DFE1D_inferenceFIM* modules), to estimate the DFE from the nonsynonymous SFSs, conditioning on the maximum-likelihood estimates of the demographic parameters. The method implemented with *fit∂a∂i* fits a DFE to the nonsynonymous data SFS by maximizing the Poisson likelihood (42). Because the Poisson likelihood requires a mutation rate for nonsynonymous variants (*θ_NS_*), we multiplied the estimates obtained for *θ_S_* by the ratio of nonsynonymous-to-synonymous mutation we estimated for the dog genome (2.21:1) and later also for the one calculated for the human genome (2.31:1 (22)). In doing so, we sought to assess whether the DFE estimates were sensitive to misspecification of this ratio, considering a plausible value for mammals.

We focused on the deleterious DFE, with selection coefficients (*s*) ranging from |*s*| = 10^-8^ to 0.5. In doing so, we considered any portion of the DFE smaller than 10^-8^ to be effectively neutral and larger than 0.5 to be lethal and have a negligible probability of being polymorphic in sample sizes considered in our study. Dominance coefficients are assumed to be *h* = 0.5, thus implying additive fitness effects.

We investigated three distributions of selection coefficients: the standard gamma distribution for DFEs (22, 49, 79); a mixture distribution where a proportion of variants are neutral, with the remaining variants with selection coefficients following a gamma distribution (“neugamma”); and the bivariate lognormal distribution, the latter exclusively for the joint estimation of DFE for each two pairs of populations (namely AW-BC, AW-LB, and AW-PG; see below). For the neugamma distribution, the neutral mass and the gamma distribution can effectively be treated as a single integrable function. The reported *s* are scaled by the inferred ancestral population size estimated according to the demographic inference using the SFS of synonymous variants, unless otherwise noted (e.g., Fig. S13). To discretize the DFE using the estimated parameters of the distributions, we computed the cumulative probability in a given range of *s* using the *pgamma* function in R.

*varDFE* outputs the standard deviation of the maximum likelihood parameter estimates (MLEs) based on the Fisher’s Information Matrix method implemented in ∂a∂i, which we then used to calculate confidence intervals using R. To do so, we assumed the MLEs of the shape and scale parameters of the gamma distribution were normally distributed with means equal to the maximum likelihood values and standard deviation as computed by *varDFE*. We then drew 10,000 shape and scale parameters using the function *pnorm* in R and used *pgamma* to compute the discretized DFE for each replicate. The 95% confidence interval for each bin was taken as the middle 95% of the simulated values.

We note that the maximum likelihood estimates for the scale parameter of the gamma distribution reached the upper boundary during the DFE inferences in some instances (see Tables S2 and S3), with uncertain biological significance. We initially used a range for the scale parameter based on previous works with different organisms and slowly increased the upper bound of the scale parameter until we reached 1,000,000. Reaching this upper bound may be due to an innately highly deleterious DFE in canids. Importantly, the fit of the model SFS to the data is satisfactory using this upper bound (Fig. S8-S11). The same phenomenon was observed by Lin et al. (56) in an independent analysis of the AW data used in this study, as well as for a population of Russian Karelian gray wolf not analyzed here. In addition, Gaughran (80) describes the same phenomenon for the DFE estimates of the northern elephant seal and the Baltic ringed seal.

### 2D Demographic and DFE Inferences

We inferred the joint DFE of pairs of populations: AW-BC, AW-LB, and AW-PG. We assume the wolf and the breed dog populations recently split form one another, and that there is symmetric gene flow between them. We also assume that the wolf population is more similar to the ancestral population and keeps the same selection coefficients while the diverged dog population may have different selection coefficients.

We inferred the demographic history and joint DFE using the joint SFS, which is a matrix in which each entry is the count of the number of variants observed at frequency *i* in population 1 and *j* in population 2. Similar to the 1D analysis, here we also used the folded allele frequency spectrum that counts the minor allele frequency. Different demographic histories and different combinations of selection coefficients of two populations lead to distinct patterns in the joint SFS (23).

We infer the demographic model using the joint SFS from synonymous variants. We assume the ancestral population split into a wolf (AW) and a breed dog (BC, LB, or PG) population. The derived populations after the split may have different population sizes among themselves and relative to the ancestral population. Gene flow is assumed to be symmetric.

Since the DFE for single populations were found to be similar, we fit the joint DFE with a symmetric bivariate lognormal model that is a joint distribution of correlated lognormal variables (mean *μ* and standard deviation *σ*) with identical marginal distributions. That is, the marginal DFE for the wolf population and dog population are the same, while the parameter *ρ* quantifies the correlation of fitness effects of mutations between populations.

We used *∂a∂i/fit∂a∂i* and the command line tool dadi-cli to infer the 2D demographic model and the joint DFE. The parameters and commands used for inference can be found at https://github.com/chenludi/canids_2DDFE_2024/tree/main.

### Gene Set Data

The AmiGO Gene Ontology (GO) database (https://amigo.geneontology.org/amigo) was accessed on 09/12/2024 for the retrieval of information regarding gene names included in GO terms putatively associated with domestication. These include “nervous system development” (GO:0007399), “immune system process” (GO:0002376), “carbohydrate metabolic process” (GO:0005975), “pigmentation” (GO:0043473), and “skeletal system development” (GO:0001501). The choice for those specific five GO terms sought to balance the specificity of the term with the number of genes included in each. The data was filtered considering *“Canis lupus familiaris*” as “Organism.” From these, we obtained the relevant gene IDs based on the “gene/product (bioentity label)” column. The coordinates for the selected genes were obtained from the UCSC Genome Browser (https://hgdownload.soe.ucsc.edu/goldenPath/canFam3/bigZips/genes/canFam3.ncbiRefSeq.g <u>tf.gz</u>), accessed on 09/05/2024. Exon length for each gene set was 5,761,394 bp, 3,703,799 bp, 1,060,844 bp, 239,818 bp, and 1,254,685 bp respectively. We considered “nervous system development” and “immune system process” separately, and also merged all five gene sets into a “domestication-associated genes” set for a total of 10,483,822 bp. Data processing and subsetting were performed with in-house scripts, as well as *BCFtools* (81) and *BEDtools* (82).

### A Recombination Map for the Arctic Wolf

To infer a fine-scale recombination map for the AW, we used the unphased high coverage polymorphism data used for the DFE inference as described above. The map was built based on linkage disequilibrium (LD) patterns across the genome. To do so, we first inferred AW demographic history (i.e., changes in *N_e_* through time) with SMC++ (83), considering all 38 autosomal chromosomes. We considered a mutation rate of 4.5e-9 per base pair per generation, based on a wolf pedigree study (84). Then, we estimated the recombination rate per-chromosome using *pyrho* (85). When computing a lookup table in *pyrho*, we used the manual recommendation of calculating statistics of LD and *rho* (population recombination rate) based on a population size that was 50% larger than our sample size. For the final step of inferring recombination rates (*r*) in *pyrho*, we used a window size and block penalty of 50.

### Assessing DFE Variation as a Function of Recombination Rate

We used the LD-based recombination map that we inferred for the AW to split its genome into three different bins based on *r* in units of per bp per generation and considering non-overlapping 1 MB windows: Low (0 ≤ *r* < 1.9e-9), Moderate (1.9e-9 ≤ *r* < 4.3e-9), and High (*r* ≥ 4.3e-9) recombination rate. We excluded regions with recombination rates above 2e-8 per bp per generation. In doing so, we sought to have approximately equal amounts of data (in base pairs) across the three bins. As a result, each bin contained roughly ⅓ the number of synonymous and nonsynonymous SNPs found across all exons. Considering each of the three recombination bins separately, we inferred demographic history and the gamma DFE using the same approach implemented for the whole exome data.

## Acknowledgments

This work is supported by a National Institutes of Health (NIH) grant R35GM119856 to KEL. CEGA was supported by the National Institute of General Medical Sciences of the NIH under award number R35GM142939. The content of this paper is solely the responsibility of the authors and does not necessarily represent the official views of the NIH. ML was supported by the David H. Smith Postdoctoral Fellowship. We thank Tanya Phung for her assistance with bioinformatics processing at an early stage of this research and contributing data, Miguel Guardado for his assistance with simulations at the initial stage of this research, and Annabel Beichman, as well as members of the Lohmueller Lab (University of California, Los Angeles) and the Malaspinas Lab (University of Lausanne) for helpful discussions.

