## Supplemental Figures for "Evolutionary consequences of domestication on the selective effects of new amino acid changing mutations in canids"

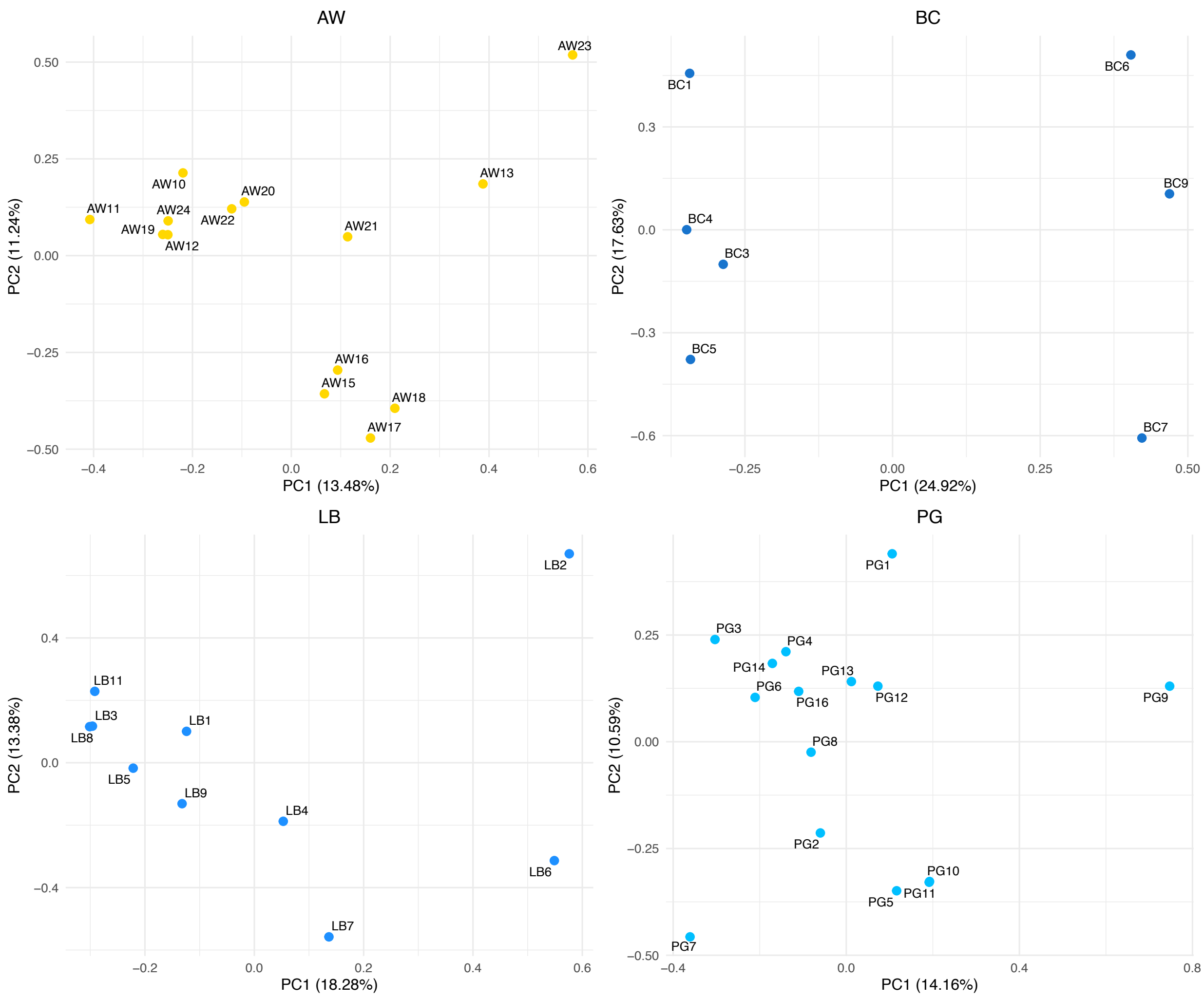


Fig. S1. Principal components analysis of population structure. The first two principal components of the data are plotted for AW, BC, LB, and PG.

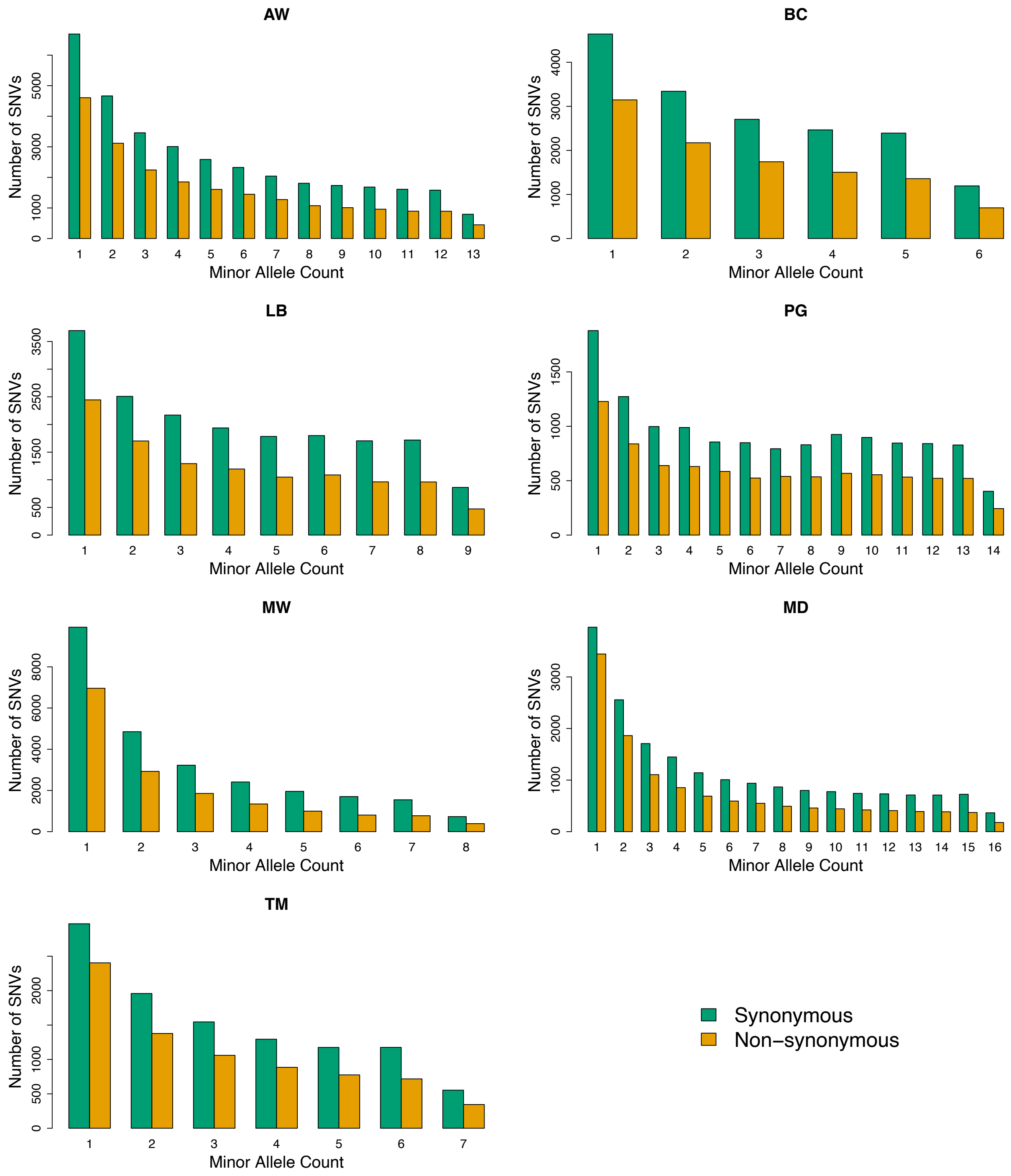


Fig. S2. The projected folded site frequency spectra (SFSs) showing the number of variants in each of the seven populations analyzed in this study. The seven populations are: AW = Arctic wolf; BC = border collie; LB = labrador retriever; PG = pug; MW = gray wolves from different populations; MD = dogs from 20 different breeds; TM = Tibetan mastiff. The teal bars represent synonymous variants, while the orange bars represent amino acid changing (nonsynonymous) variants.

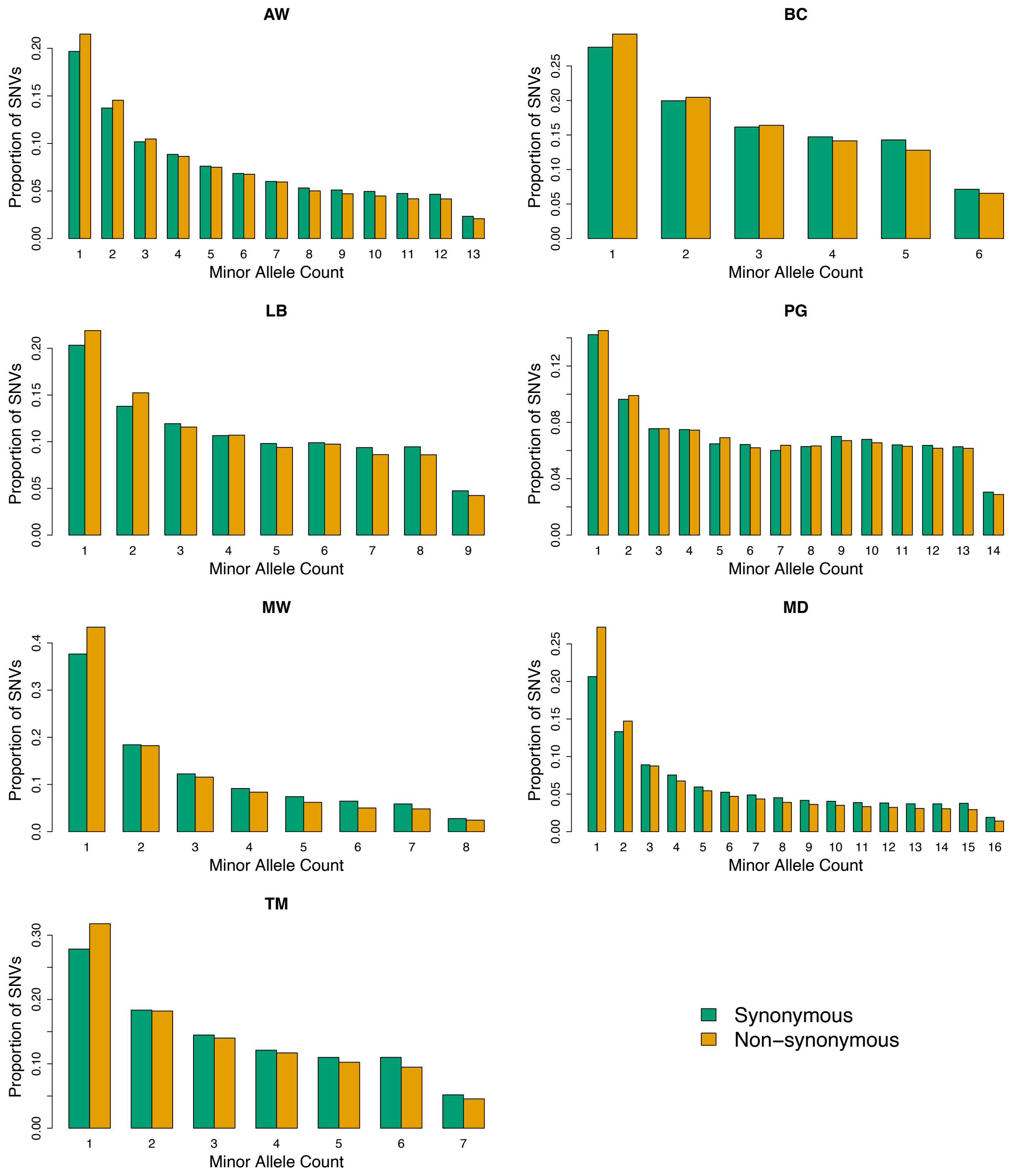


Fig. S3. The projected folded site frequency spectra (SFSs) showing the proportions of variants in each of the seven populations analyzed in this study. The seven populations are: AW = Arctic wolf; BC = border collie; LB = labrador retriever; PG = pug; MW = gray wolves from different populations; MD = dogs from 20 different breeds; TM = Tibetan mastiff. The teal bars represent synonymous variants, while the orange bars represent amino acid changing (nonsynonymous) variants.

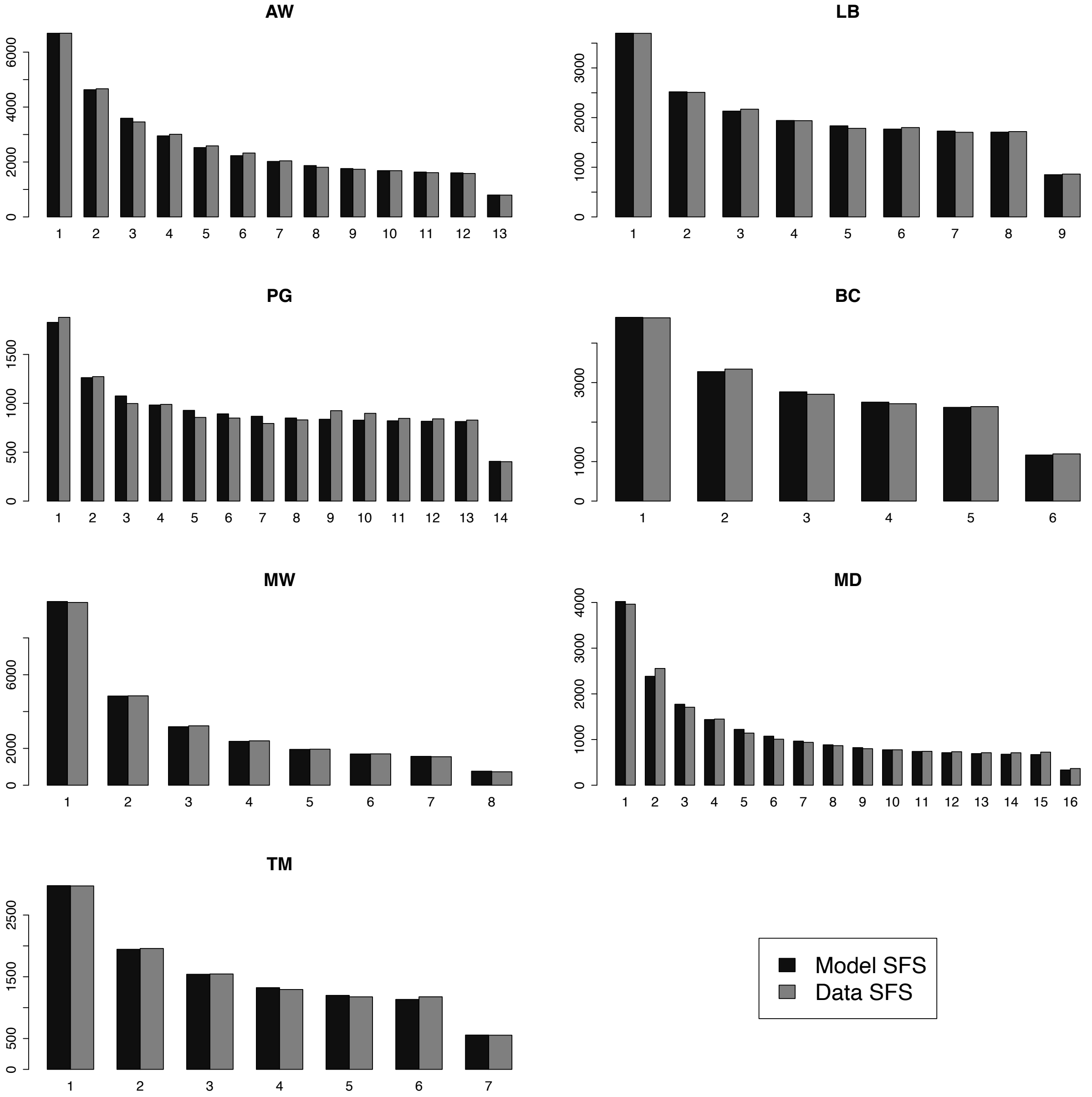


Fig. S4. Comparison of the model SFS (black) with the data synonymous SFS (gray) for the seven studied canid populations. The maximum sample size after projection is shown. The model SFS was computed based on the maximum likelihood parameters of the 2-epoch demographic models inferred in each case (Table S1).

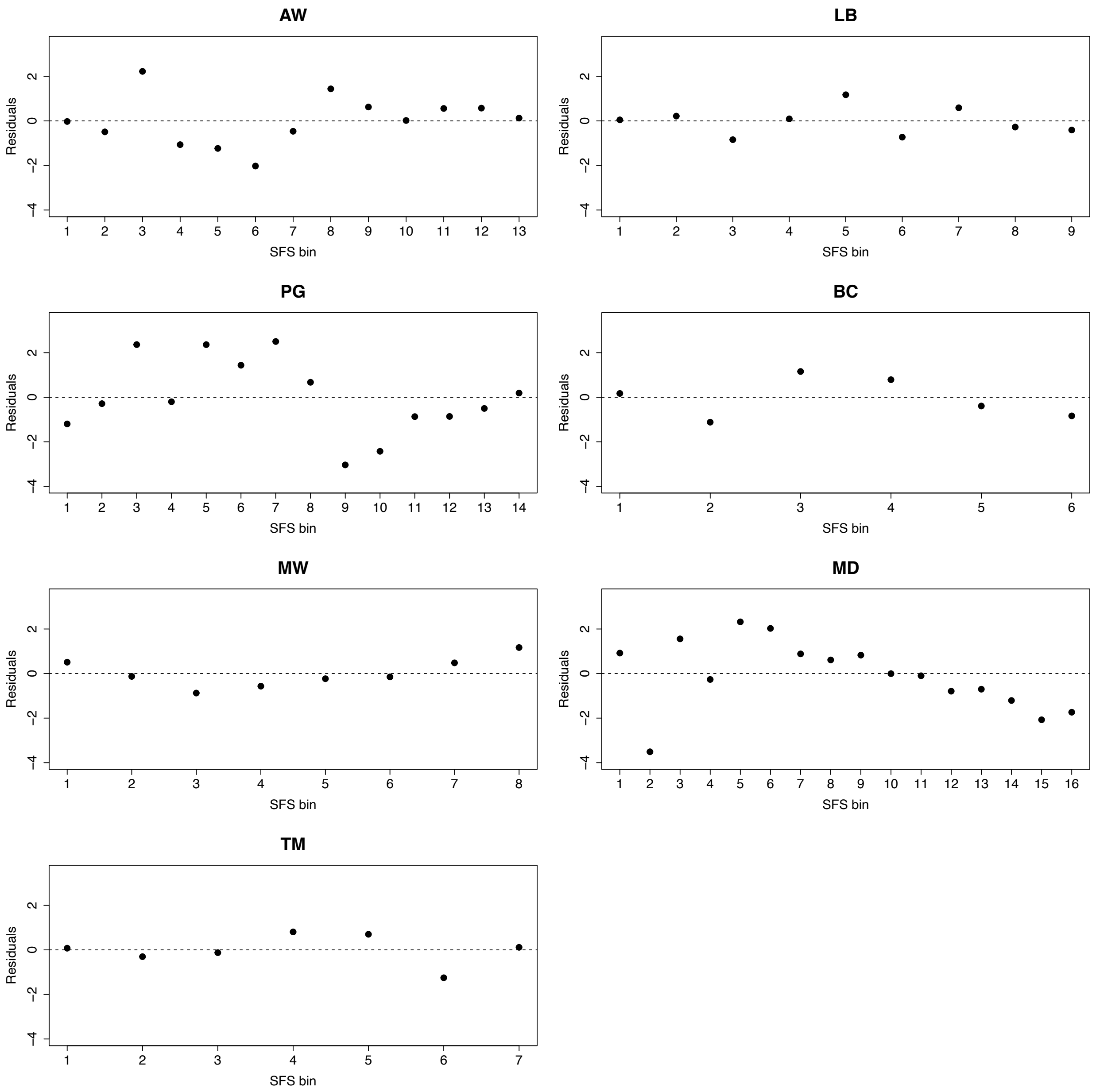


Fig. S5. Residuals of the model fit considering the model SFS (black in Fig. S4) and the observed (data) SFS (gray in Fig. S4) for the seven studied canid populations. The x-axis shows the different minor allele counts for each population. The standardized residuals (y-axis) were calculated based on the difference between the observed and the model counts in each bin, divided by the square root of the model counts.

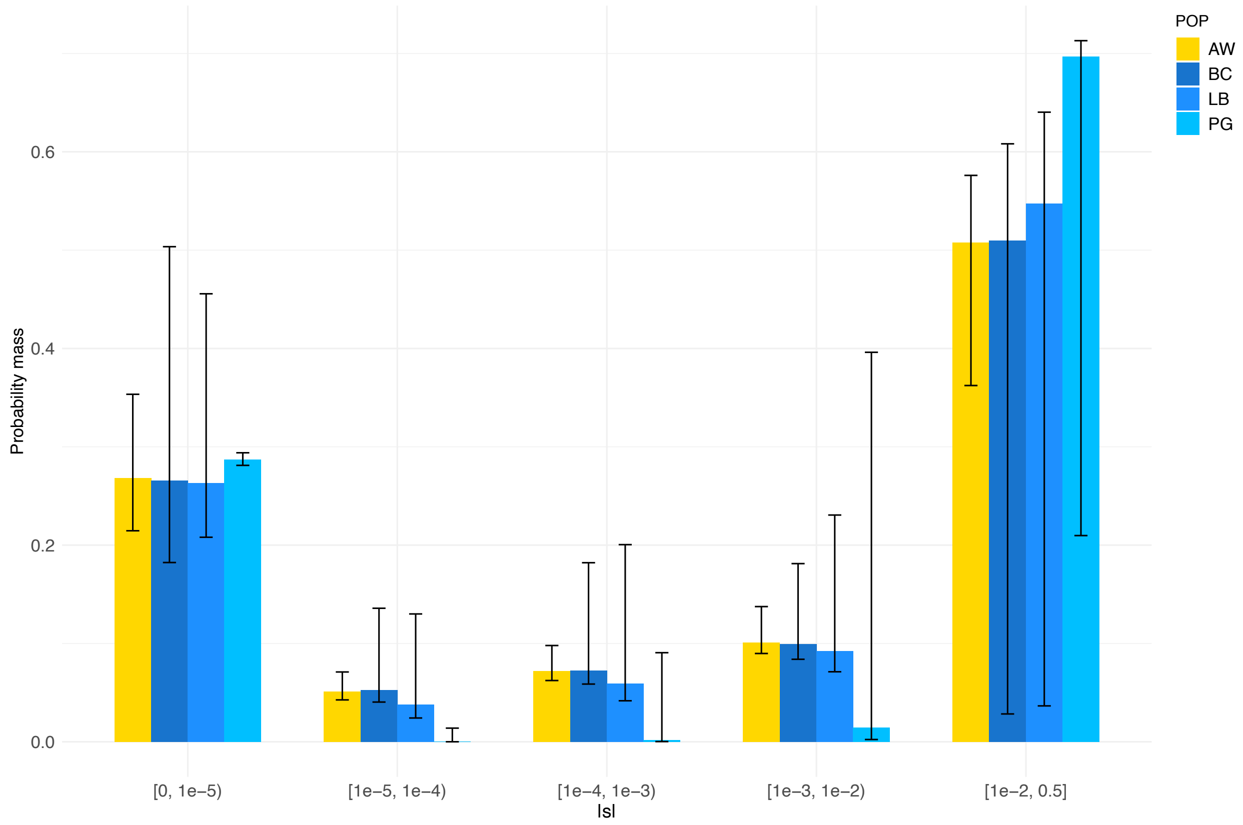


Fig. S6. Discretized distribution of fitness effects (DFE) showing the proportions of nonsynonymous mutations in various categories of $|s|$, following a neugamma distribution. From left to right, mutations range from neutral/nearly neutral (1e-5 < $|s|$ ≤ 0) to strongly deleterious/lethal ($|s| \geq$ 1e-2). For comparison, see Figure 1 for the gamma-distributed DFE. The arctic wolf population (AW) is depicted in yellow, and three domestic dog breeds are shown in different shades of blue (BC = border collie; LB = labrador retriever; PG = pug). Error bars represent the 95% confidence intervals for the proportion of mutations in each category of $|s|$.

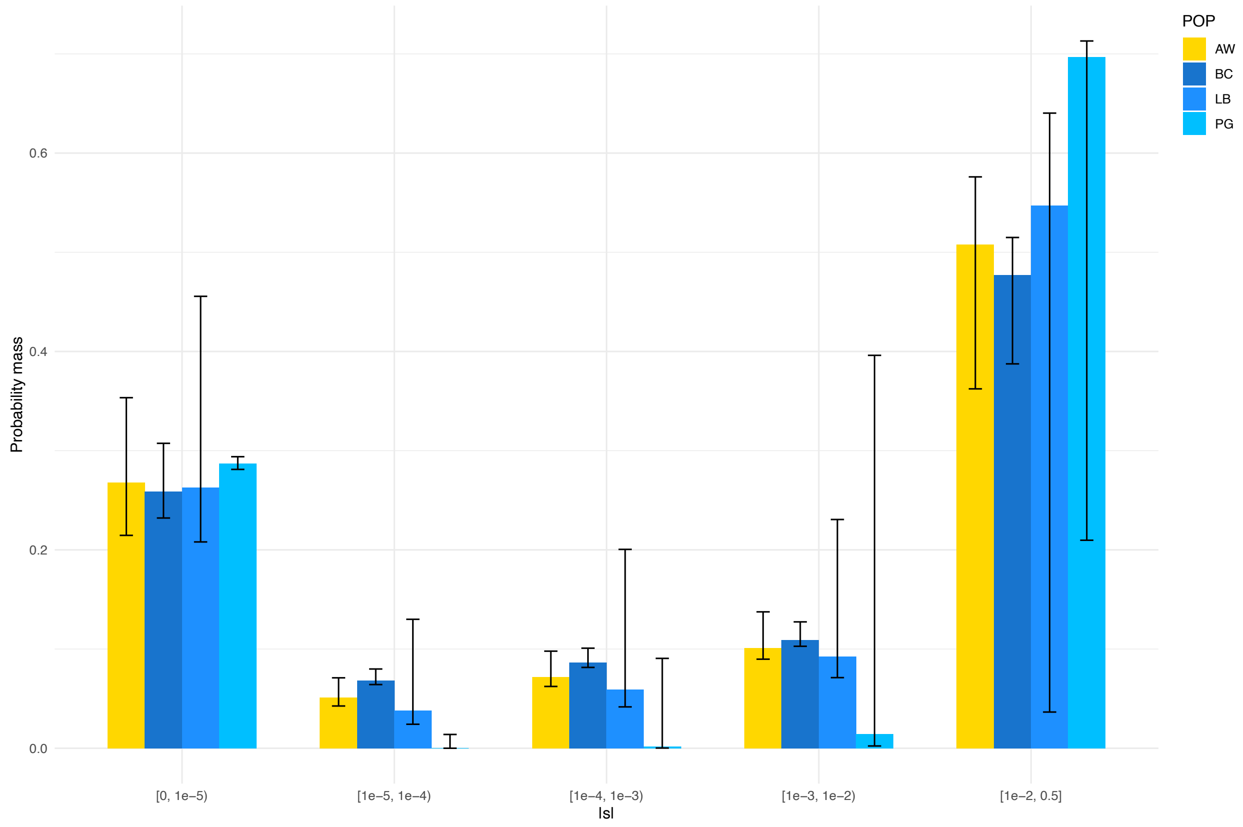


Fig. S7. Discretized distribution of fitness effects (DFE) showing the proportions of nonsynonymous mutations in various categories of $|s|$. From left to right, mutations range from neutral/nearly neutral ($1e\text{-}5 < |s| \leq 0$) to strongly deleterious/lethal ($|s| \geq 1e\text{-}2$). Shown are the results for the neugamma DFE for AW, LB, and PG (same data as Fig. S6) and the gamma DFE for BC (same data as Fig. 1). The neugamma DFE improved the fit for AW, LB, and PG relative to the gamma DFE, although only marginally so. Conversely, the gamma DFE presented a better fit relative to the neugamma DFE for BC. The arctic wolf population (AW) is depicted in yellow, and the three domestic dog breeds are in different shades of blue (BC = border collie; LB = labrador retriever; PG = pug). Error bars represent the 95% confidence intervals for the proportion of mutations in each category of $|s|$.

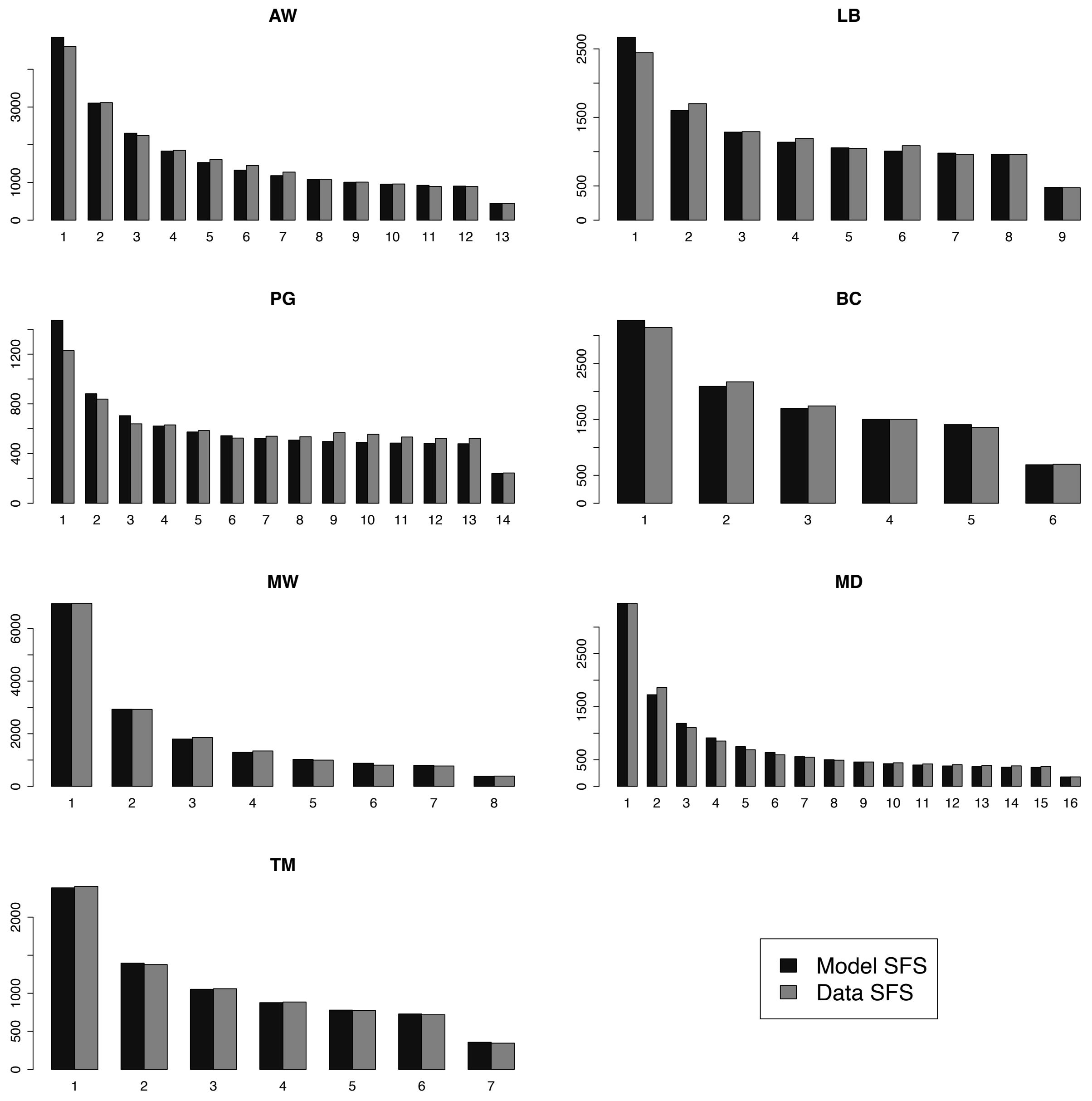


Fig. S8. Comparison of the model SFS (black) with the data nonsynonymous SFS (gray) for the seven studied canid populations. The maximum sample size after projection is shown. The model SFS was computed based on the maximum likelihood parameters of the gamma-distributed DFE (Table S2).

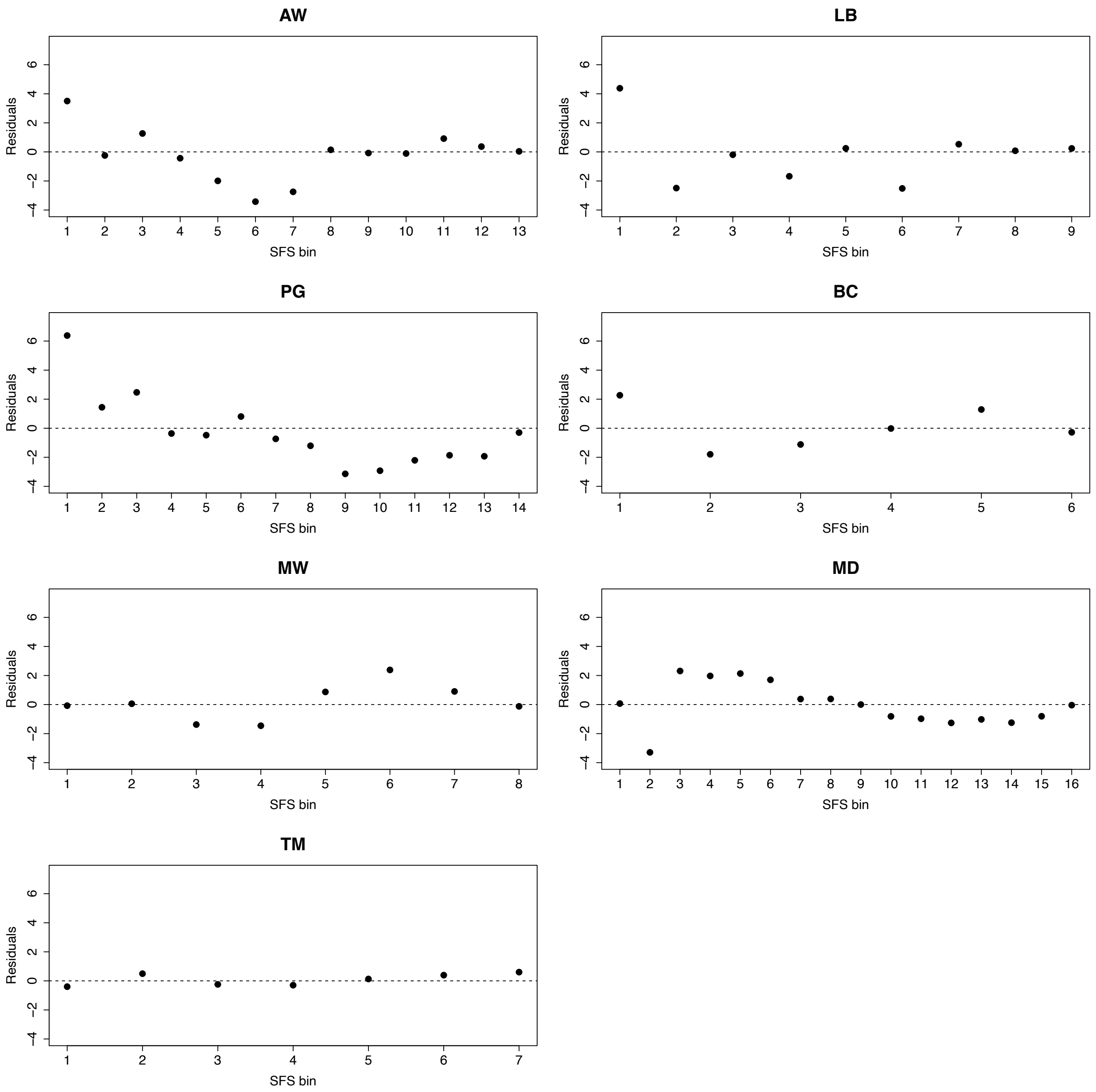


Fig. S9. Residuals of the model fit considering the model SFS (black in Fig. S8) and the observed (data) SFS (gray in Fig. S8) for the seven studied canid populations. The x-axis shows the different minor allele counts for each population. The standardized residuals (y-axis) were calculated based on the difference between the observed and the model counts in each bin, divided by the square root of the model counts.

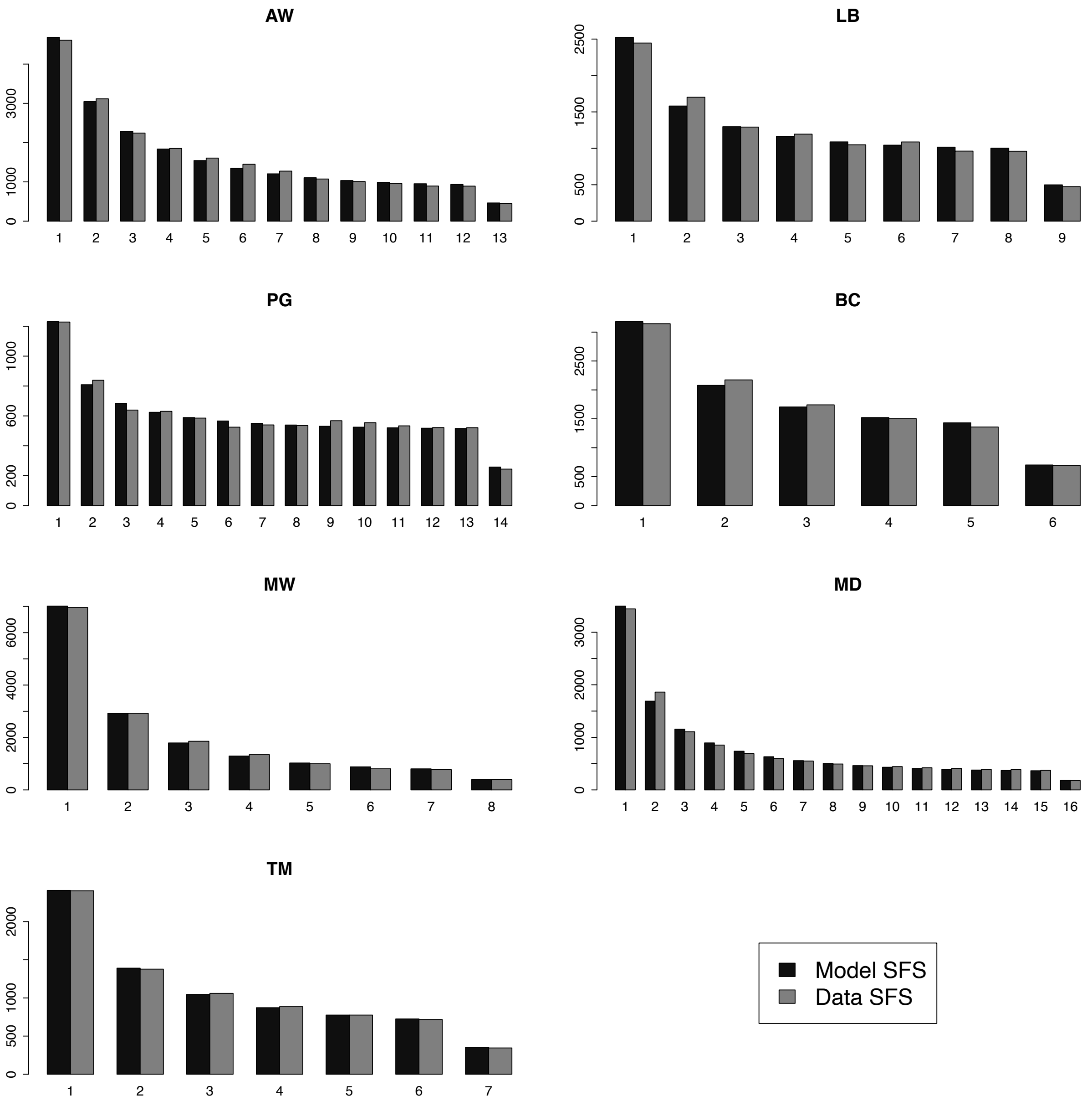


Fig. S10. Comparison of the model SFS (black) with the data nonsynonymous SFS (gray) for the seven studied canid populations. The maximum sample size after projection is shown. The model SFS was computed based on the maximum likelihood parameters considering the neugamma distribution for the DFE (Table S3).

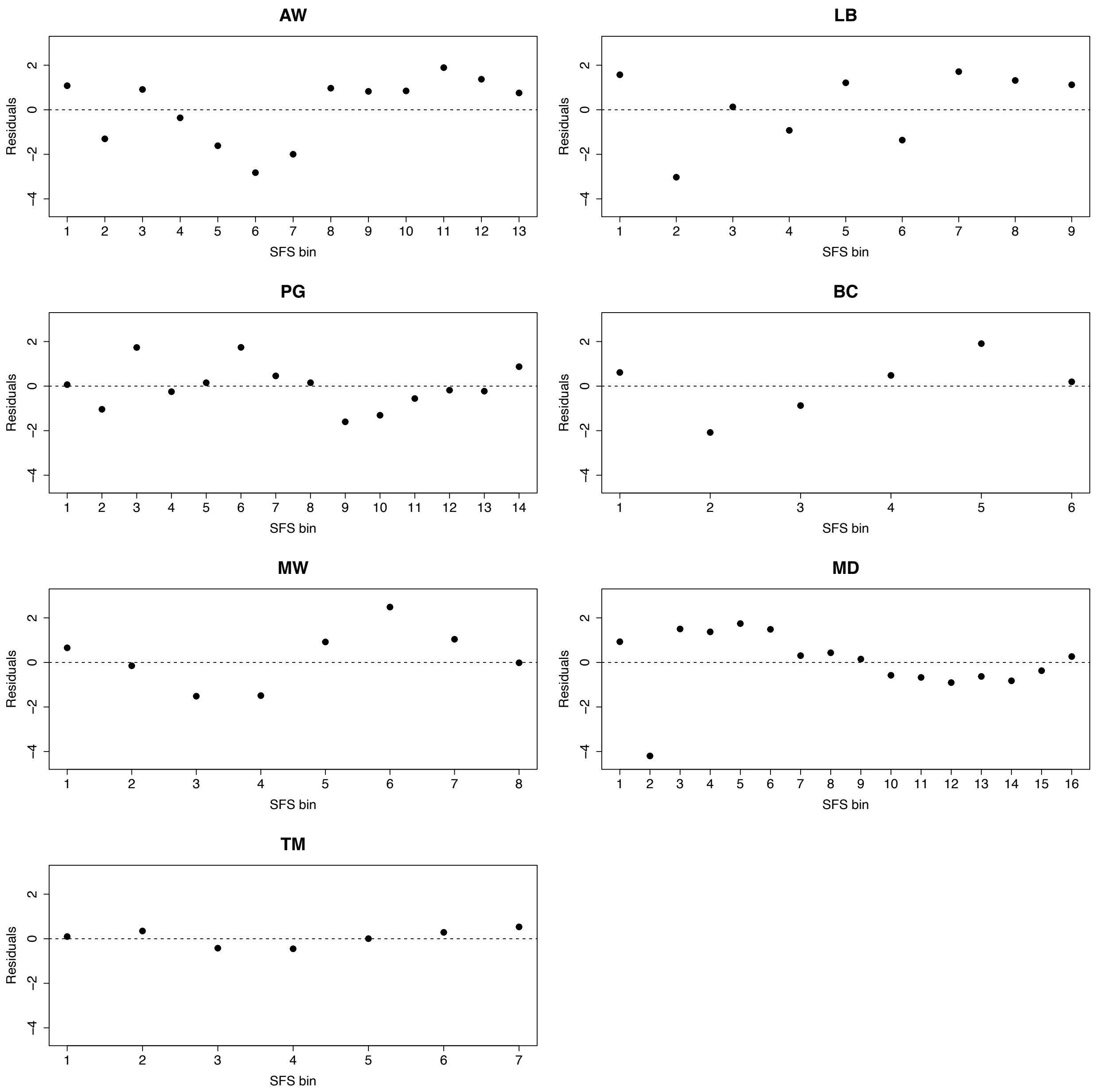


Fig. S11. Residuals of the model fit considering the model SFS (black in Fig. S10) and the observed (data) SFS (gray in Fig. S10) for the seven studied canid populations. The x-axis shows the different minor allele counts for each population. The standardized residuals (y-axis) were calculated based on the difference between the observed and the model counts in each bin, divided by the square root of the model counts.

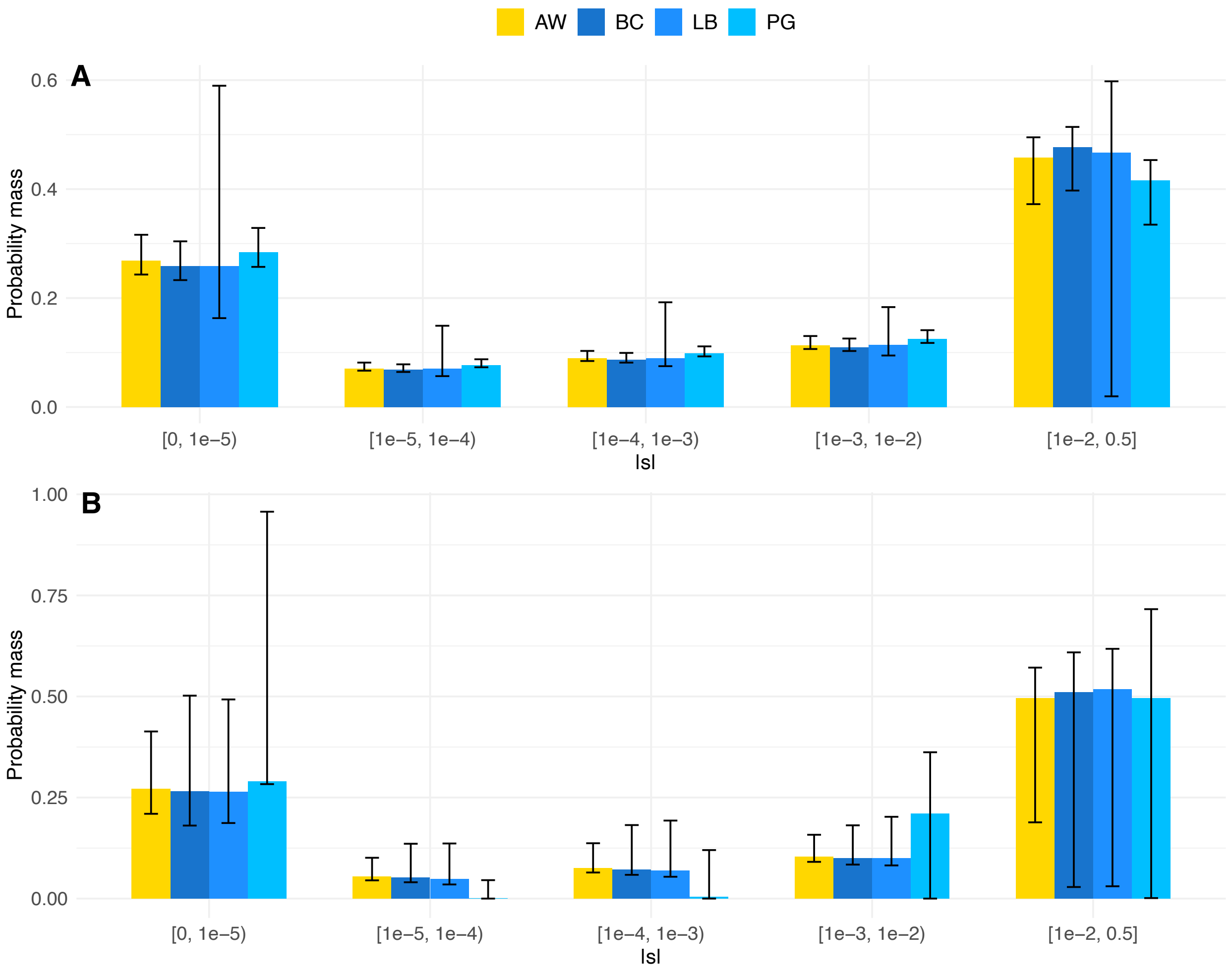


Fig. S12: Discretized DFE estimates using equal sample sizes ($n_{eq}$ = 6) across populations (AW = Arctic wolf; BC = border collie; LB = labrador retriever; PG = pug) assuming (A) a gamma and (B) neugamma distribution for $|s|$.

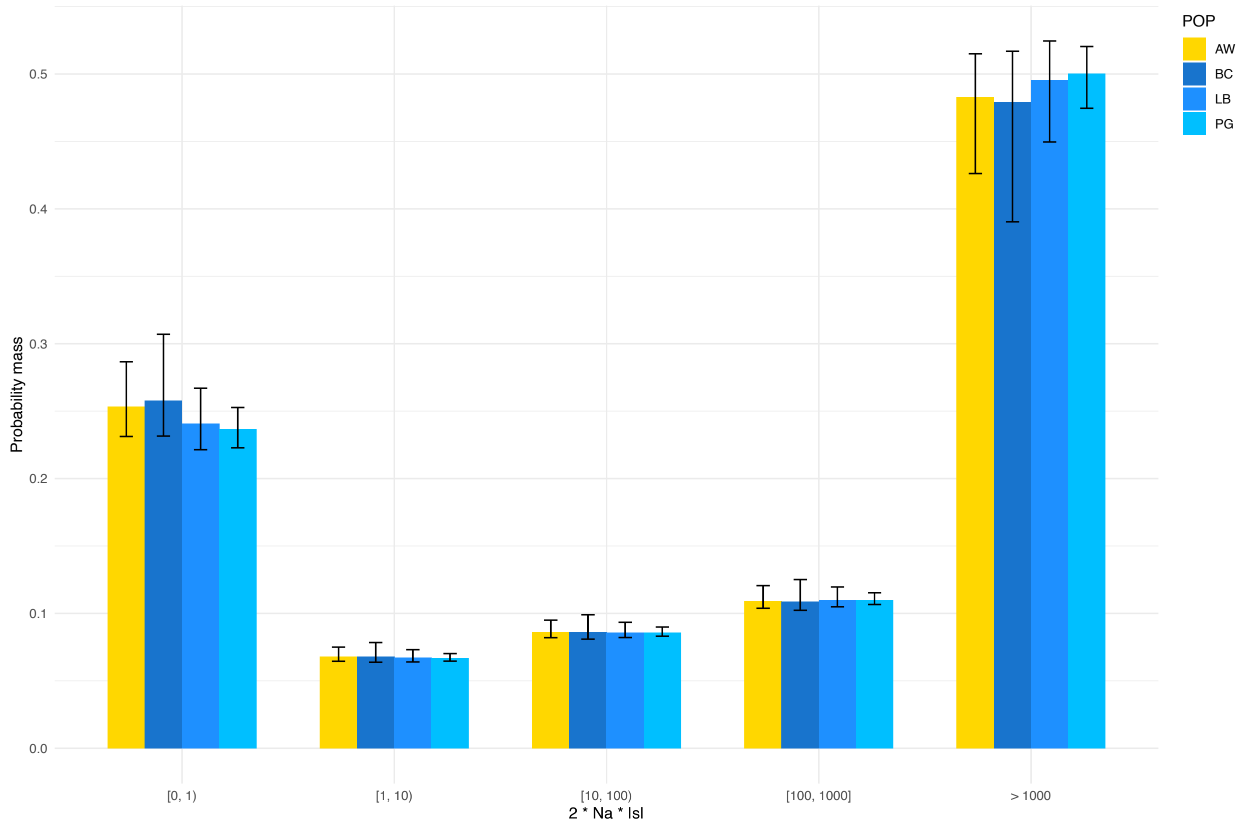


Fig. S13. Discretized distribution of fitness effects (DFE) in terms of $2N_a s$. From left to right, mutations range from neutral/nearly neutral ($2N_a s < 1$) to strongly deleterious/lethal ($2N_a s > 1000$). The DFE is assumed to follow a gamma distribution. The arctic wolf population (AW) is depicted in yellow, and the three domestic dog breeds are shown in different shades of blue (BC = border collie; LB = labrador retriever; PG = pug). Error bars represent the 95% confidence intervals for the proportion of mutations in each category of $|s|$.

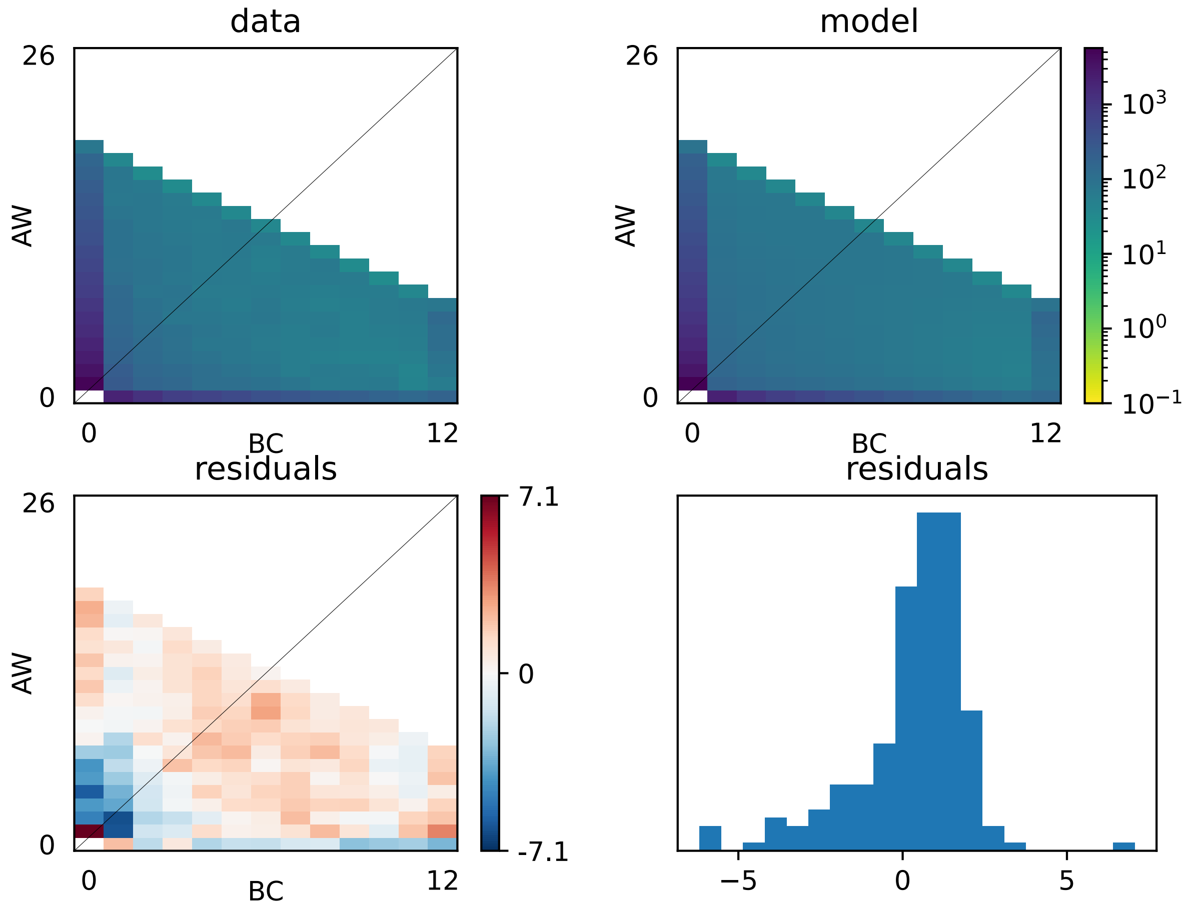


Fig. S14. Joint synonymous SFS for the Arctic wolf (AW) and the border collie (BC). The upper panels show the joint 2D synonymous SFSs for the data (top left) and the model (top right) considering the "split_mig" demographic model in *∂a∂i*. The bottom panels show the residuals between the model and the data. Specifically, the bottom left panel shows the residuals per SFS cell, and the bottom right the distribution of the residuals.

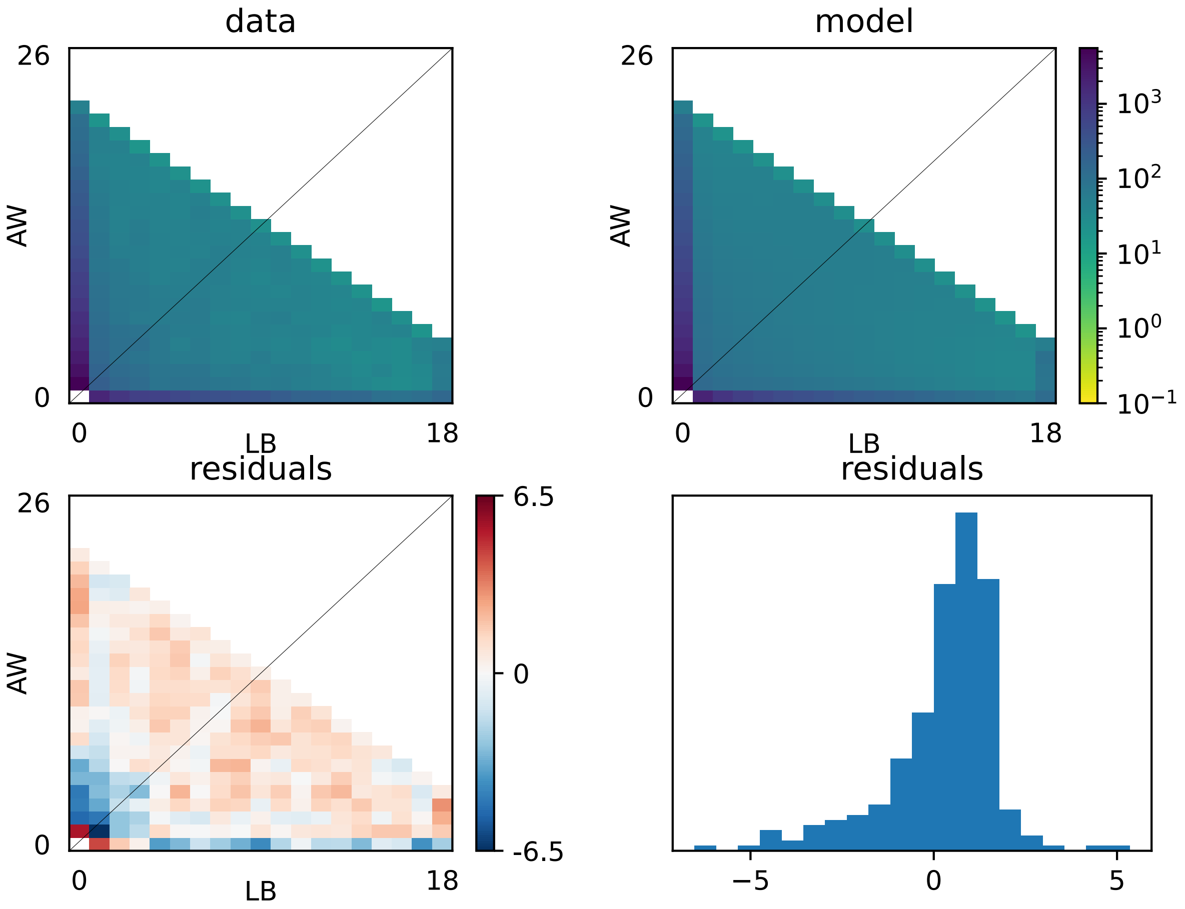


Fig. S15. Joint synonymous SFS for the Arctic wolf (AW) and the labrador retriever (LB). The upper panels show the joint 2D synonymous SFSs for the data (top left) and the model (top right) considering the "split_mig" demographic model in *∂a∂i*. The bottom panels show the residuals between the model and the data. Specifically, the bottom left panel shows the residuals per SFS cell, and the bottom right the distribution of the residuals.

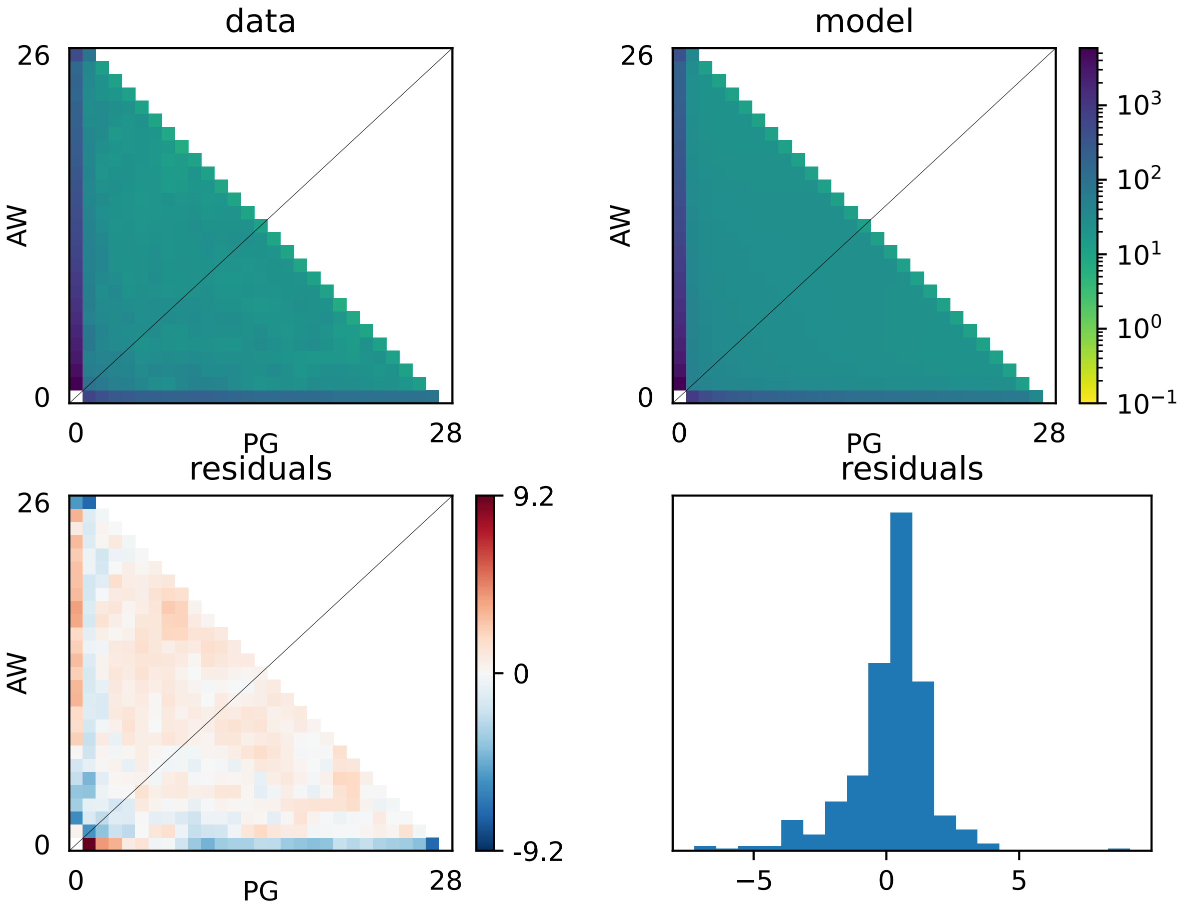


Fig. S16. Joint synonymous SFS for the Arctic wolf (AW) and the pug (PG). The upper panels show the joint 2D synonymous SFSs for the data (top left) and the model (top right) considering the "split_mig" demographic model in *∂a∂i*. The bottom panels show the residuals between the model and the data. Specifically, the bottom left panel shows the residuals per SFS cell, and the bottom right the distribution of the residuals.

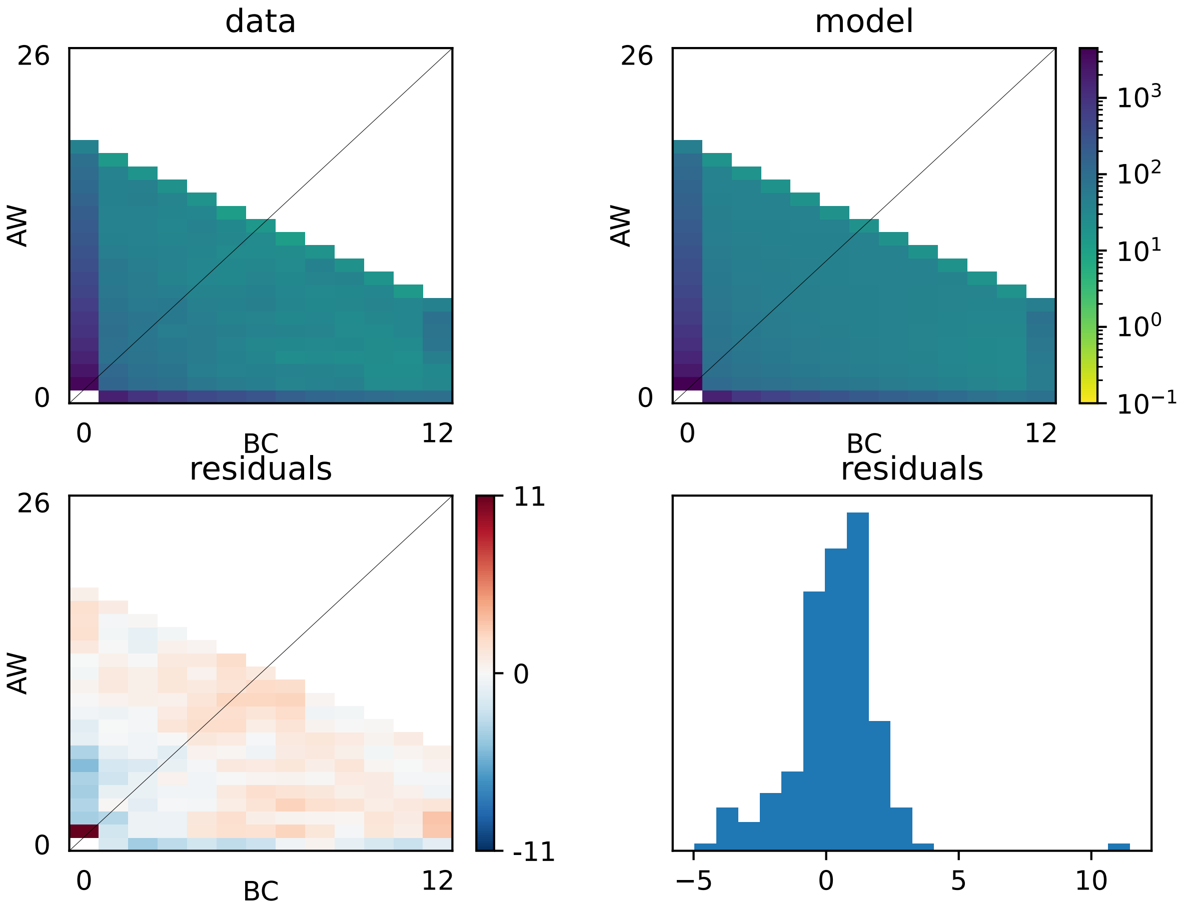


Fig. S17. Joint nonsynonymous SFS for the Arctic wolf (AW) and the border collie (BC). The upper panels show the joint 2D synonymous SFSs for the data (top left) and the model (top right) considering a bivariate lognormal distribution of $|s|$. The bottom panels show the residuals between the model and the data. Specifically, the bottom left panel shows the residuals per SFS cell, and the bottom right the distribution of the residuals.

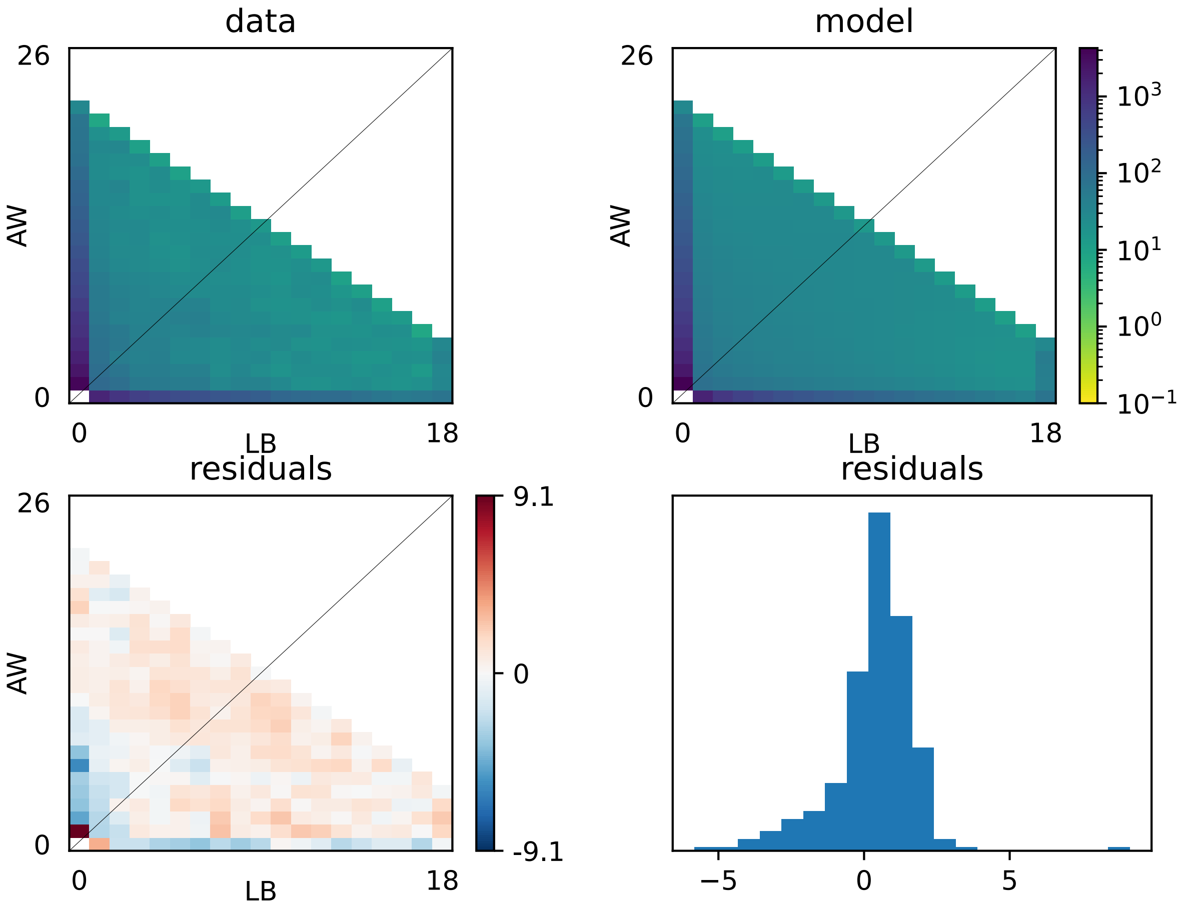


Fig. S18. Joint nonsynonymous SFS for the Arctic wolf (AW) and the labrador retriever (LB). The upper panels show the joint 2D nonsynonymous SFSs for the data (top left) and the model (top right) considering a bivariate lognormal distribution of $|s|$. The bottom panels show the residuals between the model and the data. Specifically, the bottom left panel shows the residuals per SFS cell, and the bottom right the distribution of the residuals.

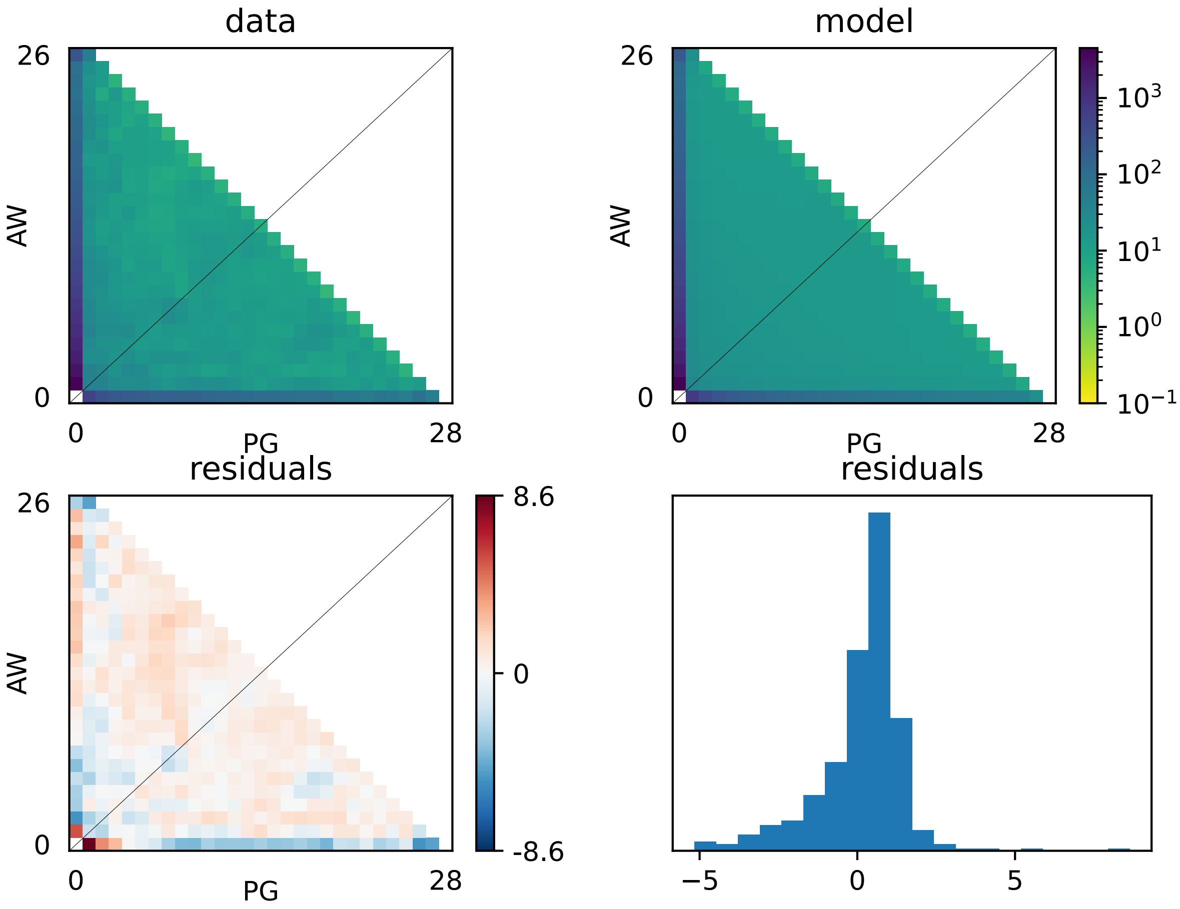


Fig. S19. Joint nonsynonymous SFS for the Arctic wolf (AW) and the pug (PG). The upper panels show the joint 2D nonsynonymous SFSs for the data (top left) and the model (top right) considering a bivariate lognormal distribution of $|s|$. The bottom panels show the residuals between the model and the data. Specifically, the bottom left panel shows the residuals per SFS cell, and the bottom right the distribution of the residuals.

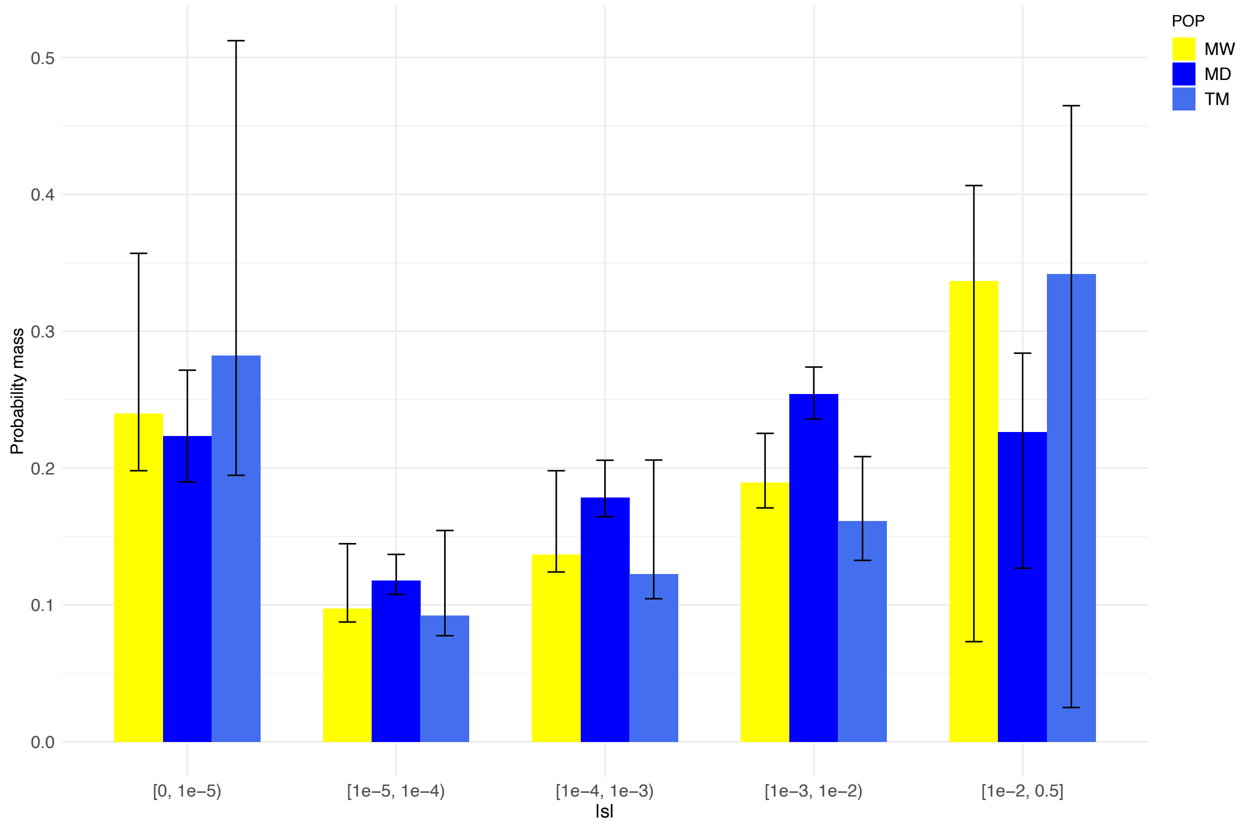


Fig. S20. Discretized distribution of fitness effects (DFE) showing the proportions of nonsynonymous mutations in various categories of $|s|$. From left to right, mutations range from neutral/nearly neutral ($0 < |s| \leq 1e\text{-}5$) to strongly deleterious/lethal ($|s| \geq 1e\text{-}2$). The DFE is assumed to follow a gamma distribution. Wolves (MW) are depicted in yellow and breed dogs (MD and TM) in different shades of blue. Error bars represent the 95% confidence intervals for the proportion of mutations in each category of $|s|$.

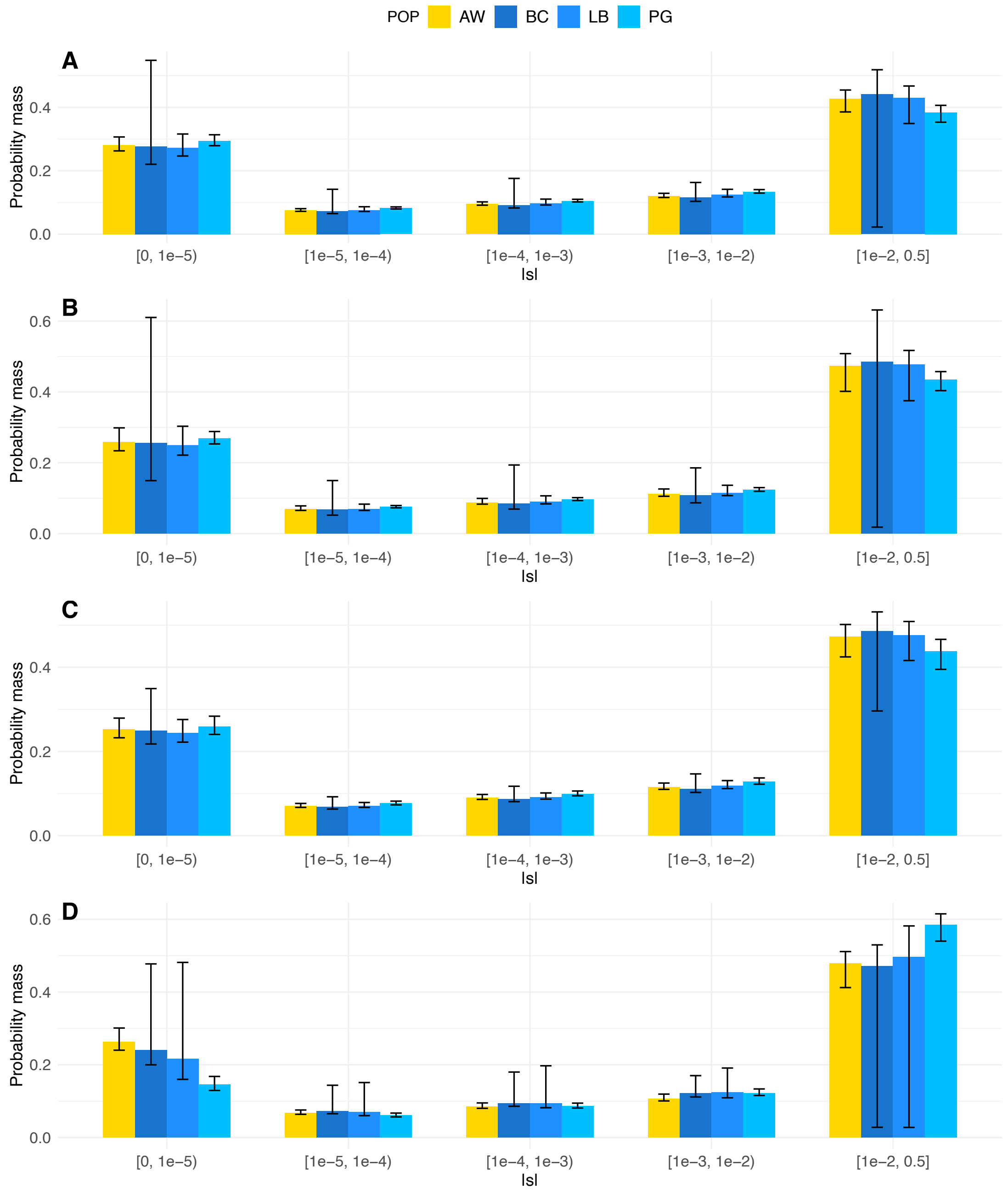


Fig. S21. Discretized distribution of fitness effects (DFE) showing the proportions of nonsynonymous mutations in various categories of $|s|$ according to different assumptions: (A) a low mutation rate of 3.00e-9 per base pair per generation, (B) a high mutation rate of 6.73e-9 per base pair per generation, (C) the human expected NS:S ratio of 2.31, and (D) a 3-epoch demographic model. In all three panels, mutations range from neutral/nearly neutral (1e-5 $< |s| \leq$ 0) to strongly deleterious/lethal ($|s| \geq$ 1e-2) and the DFE is assumed to follow a gamma distribution. The arctic wolf population (AW) is depicted in yellow, and the three different domestic dog breeds are shown in different shades of blue (BC = border collie; LB = labrador retriever; PG = pug). Error bars show the 95% confidence intervals.

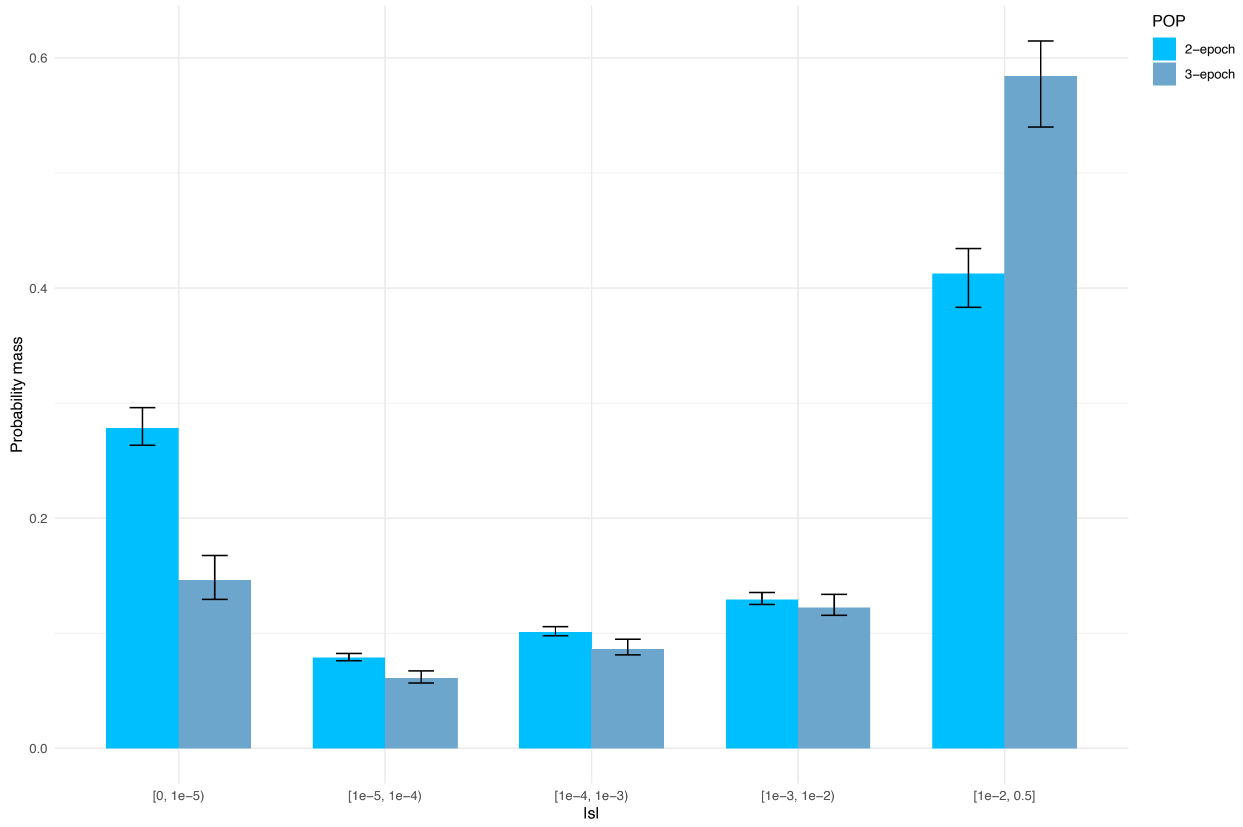


Fig. S22. Comparison of the discretized gamma-distributed DFE for pug assuming 2-epoch and 3-epoch demographic models. The DFE shows the proportions of nonsynonymous mutations in various categories of $|s|$. From left to right, mutations range from neutral/nearly neutral (1e-5 < $|s|$ ≤ 0) to strongly deleterious/lethal ($|s|$ ≥ 1e-2). Error bars represent the 95% confidence intervals for the proportion of mutations in each category of $|s|$.

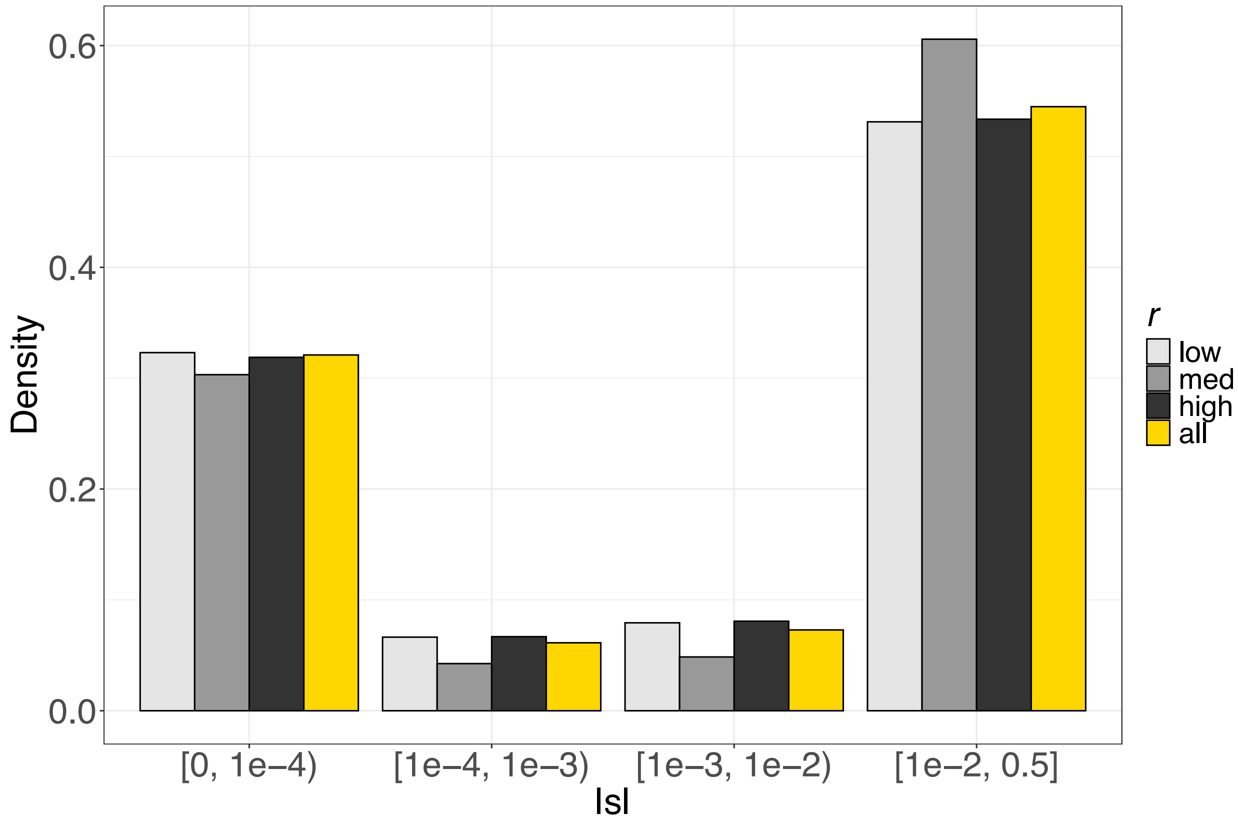


Fig. S23. Comparison of the AW DFE for different recombination rates. 1 MB windows of the genome were classified into three bins with different recombination rates ($r$) in units of per bp per generation: low $r$ ($0 \leq r < 1.9e\text{-}9$), moderate $r$ ($1.9e\text{-}9 \leq r < 4.3e\text{-}9$), and high $r$ ($4.3e\text{-}9 \leq r \leq 2e\text{-}8$). DFEs for these three bins and all the exome are color-coded: low $r$ (blue), moderate $r$ (purple), high $r$ (red), and all exome (white).

### Supplemental Table Legends

Table S1. Demographic parameter estimates considering a 2-epoch model.

Table S2. Selection parameter estimates considering gamma-distributed selection coefficients.

Table S3. Selection parameter estimates considering a neugamma distribution for the DFE.

Table S4. 2D demographic and selection parameter estimates considering three pairs of populations AW-BC, AW-LB, and AW-PG.

Table S5. Demographic parameter estimates considering a 3-epoch model.

Table S6. Shown are the estimated proportion of mutations in different ranges of $|s|$ and the 95% confidence intervals considering different gene sets. Mutations range from neutral/nearly neutral ($1e\text{-}5 < |s| \leq 0$) to strongly deleterious/lethal ( $|s| \geq 1e\text{-}2$). These data were used to construct the plot in Fig. 2, showing the discretized distribution of fitness effects (DFE) for different gene sets: Nervous System Development, Immune System Processes, and a combination of Immune System Processes, Nervous System Development, Carbohydrate Metabolic Processes, Pigmentation, and Skeletal System Development ("Domestication Genes" subset).
